# Dicer promotes genome stability via the bromodomain transcriptional co-activator Brd4

**DOI:** 10.1101/2021.01.08.425946

**Authors:** MJ Gutbrod, B Roche, JI Steinberg, AA Lakhani, K Chang, AJ Schorn, RA Martienssen

## Abstract

RNA interference is essential for transcriptional silencing and genome stability, but conservation of this role in mammals has been difficult to demonstrate. *Dicer1*^-/-^ mouse embryonic stem cells have microRNA-independent proliferation defects, and we conducted a CRISPR-Cas9 screen to restore viability. We identified suppressor mutations in transcriptional activators, H3K9 methyltransferases, and chromosome segregation factors, strongly resembling Dicer suppressors in fission yeast. Suppressors rescued chromosomal defects, and reversed strand-specific transcription of major satellite repeats in *Dicer1*^-/-^. The strongest suppressors were in *Brd4*, and in the transcriptional elongator/histone acetyltransferase *Elp3*. Using viable mutants and pharmaceutical inhibitors, we demonstrate that deletion of specific residues in *Brd4* rescue genome instability defects of *Dicer1*^-/-^ in both mammalian cells and fission yeast, implicating Dicer in coordinating transcription and replication of satellite repeats.

**Summary:** Replication and segregation defects in *Dicer1*^*-/-*^ stem cells depend on centromeric transcription by *Brd4*, and are deeply conserved in fission yeast.

---

Canonical RNA interference (RNAi) silences genes and transposable elements (TEs) through microRNAs *(1)*, small interfering RNAs (siRNAs), or PIWI-interacting RNAs (piRNAs) *(2)*. However, RNAi also controls genome instability via chromosome dosage and segregation *(3–7)*, transcription termination, and the DNA damage response *(8–13)*. For example, the fission yeast *Schizosaccharomyces pombe* does not have microRNAs, but RNAi plays an important role in heterochromatic silencing and chromosome segregation through a process called co-transcriptional gene silencing (CTGS) *(14, 15)*. Chromosome segregation defects in RNAi mutants are associated with a reduction in histone H3K9 methylation at the centromere *(16, 17)*, increased transcription, and the loss of cohesin at the locus. Additionally, RNAi becomes essential in the cell divisions preceding quiescence *(18)*. Dicer (Dcr1) and CTGS remove RNA polymerases that would otherwise transcribe centromeric repeats, preventing collision with replication and DNA damage *(12, 13)*. Polymerase removal is also required for long term survival in quiescence *(18)*.

Similar mitotic chromosomal defects result from perturbing Dicer in other eukaryotes *(19–22)* including human *(21, 23)* and mouse cells *(24)*. While the deletion of the microRNA-specific factor *Dgcr8* in mouse embryonic stem cells (mESCs) generates only mild cell cycle stalling in G1 *(25)*, the deletion of *Dicer1* in mESCs generates severe proliferation defects, strong stalling in G1, a significant increase in apoptosis, and an accumulation of transcripts from the major (pericentromeric) and minor (centromeric) satellites, all of which must be microRNA-independent *(26–28)*. In the mouse, DICER1 associates with major satellite RNA and pericentromeric chromatin *(29)* which is characterized by H3K9me2/3 and HP1 proteins as in fission yeast *(30, 31)*. H3K27me3 is also present and is redundant with H3K9me2/3 *(32, 33)*. In *Dicer1*^-/-^ mESCs, widely differing phenotypes have been reported *(26–28)* and one explanation might be the accumulation of mutations that allow stalled *Dicer1*^-/-^ cells to proliferate *(28)*. Centromeric chromatin-associated genetic suppressors arise in Dicer mutants of fission yeast when they exit the cell cycle and as RNAi becomes essential, resulting in the selection and outgrowth of suppressed strains *(18)*. We hypothesized that when *Dicer1*^-/-^ mESCs stall in G1, an essential function is revealed that needs to be suppressed in a similar way. We therefore performed a CRISPR-Cas9 genetic screen in *Dicer1*^-/-^ mESCs, focused on chromatin modifiers found in similar screens of *S. pombe (18)* to identify genetic suppressors and characterize the molecular mechanism of the *Dicer1* viability defects.

## RESULTS

### *Dicer1* is Essential for Proliferation and Chromosome Segregation in Mouse ES Cells

We reproduced the severe phenotype of *Dicer1*^-/-^ mESCs by using inducible homozygous deletion of the RNase III domains upon treatment with hydroxytamoxifen (OHT) at day 0 (d0) *(28, 34)*. We also derived stable clonal lines from rare single mutant cells that survived induction *(Dicer1*^-/-^ clones), but only after several weeks, indicating selection for suppressors had occurred (Fig. S1A). During the induction timecourse, we observed strong proliferation defects, accumulation of cells in G1, and increased cell death and DNA damage (Fig. S1B,C), as previously reported *(26, 28)*. We also observed frequent chromosome lagging, bridging, and micronuclei indicating segregation defects (Figs. 1A, 1B, S2A and S2B). No such phenotypes were observed in a *Dgcr8* knockout cell line (Figs. 1B and S2B), ruling out microRNA-based mechanisms *(23)*. Whole genome sequencing revealed partial trisomy in two *Dicer1*^-/-^ clonal cell lines (Fig. 1C), in addition to aneuploidies commonly found in mESC, that might contribute to survival *(35, 36)*. Transcriptome analysis of these lines (Fig. 1D) also agreed with previous studies, including increased expression of apoptosis-promoting genes (Fig. S3A), but the most upregulated non-coding transcripts were from pericentromeric satellite repeats and endogenous retroviruses (ERVs) (Fig. 1D and Table S1), potentially accounting for the strong interferon response (Fig. 1E) *(26, 37)*. Transcriptomes from *Dicer1*^-/-^ clonal lines differed substantially from that of freshly induced cells, consistent with strong selection for viable clones (Fig. S3B). Intriguingly, we observed a reverse strand bias in major satellite transcripts in *Dicer1*^-/-^ mESCs that switched to the forward strand after selection for viable clones (Fig. 1F). No such switches were observed in other genes or TEs genome wide.

**Figure 1.**
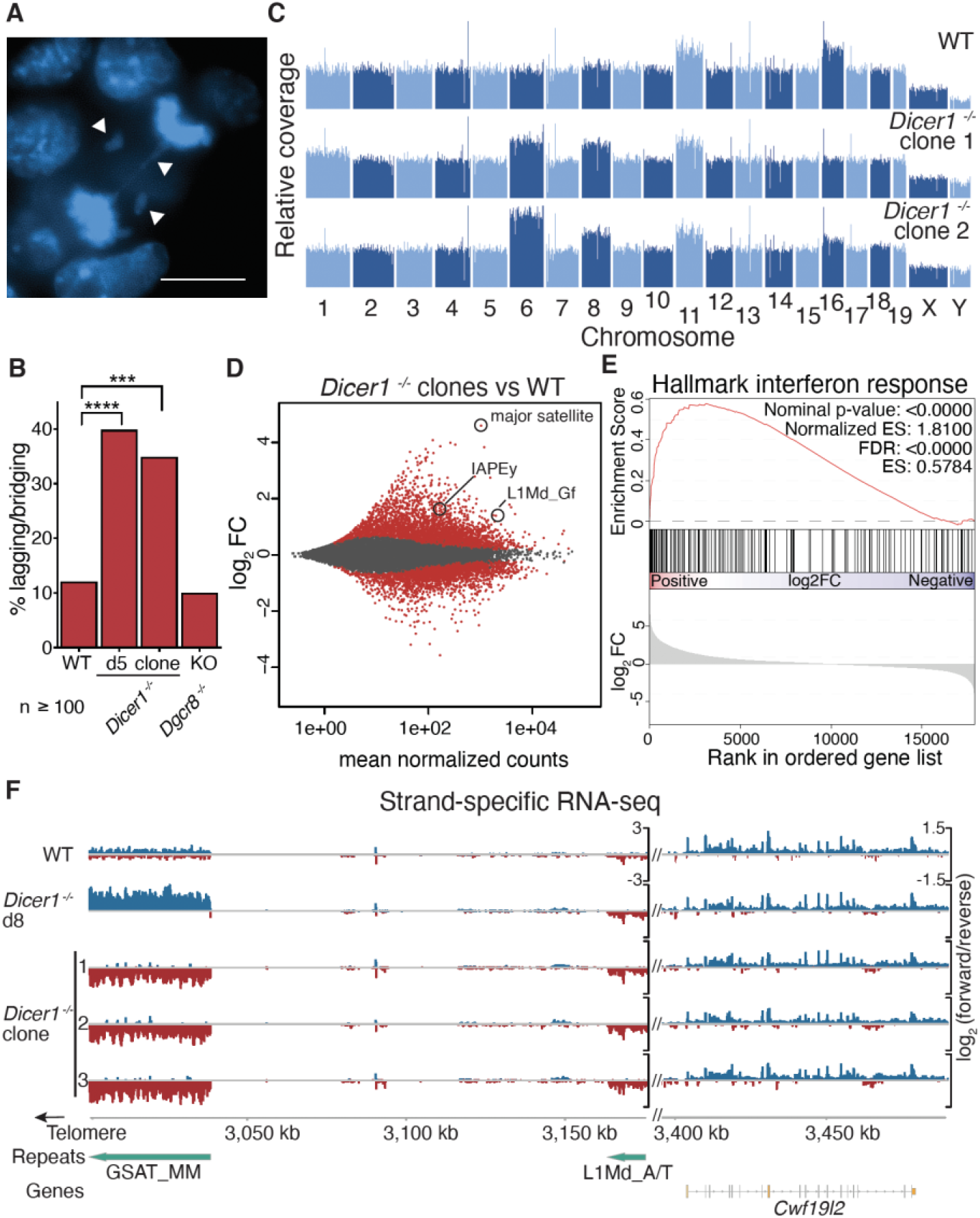
*Dicer1* promotes proliferation and chromosome segregation in mESCs and regulates the major satellite transcript in a strand-specific manner (A) DAPI staining of lagging and bridging chromosomes (arrows) in *Dicer1*^-/-^ anaphase cells. (B) Quantification of lagging/bridging chromosomes (**** - p< 0.0001, *** − 0.001 >p> 0.0001, − Fisher’s exact test). Scale bar = 10 microns (n > 100). Chromosome mis-segregation is prevalent in *Dicer1*^-/-^ cells 5 days after deletion of the *Dicer1* gene (d5) as well as in clones selected for viability after several weeks (clones), but not in cells deficient for the miRNA processor *Dcgr8*. (C) Genome browser of normalized coverage of whole genome sequencing reads reveals enhanced aneuploidy in *Dicer1*^-/-^ clones. Mouse autosomes are numbered 1-19. (D) Differential expression analysis of *Dicer1*^*-/-*^ clones compared to uninduced wild type cells by RNA-seq. The major satellite repeat is strongly upregulated. Other non-coding transcripts include endogenous retroviruses (IAP) and LINE elements (L1). (E) Hallmark Gene Sets Enrichment Analysis (GSEA) reveals an activated interferon response in *Dicer1*^-/-^ clones. (F) Strand-specific transcriptional upregulation of a 38kb major satellite repeat (GSAT_MM) on chr9, but not of nearby L1 TEs or genes. The log_2_ normalized ratio of forward (blue) and reverse (red) strand is plotted. One representative replicate of 3 is plotted for WT and *Dicer1*^*-/-*^ cells 8 days after induction (d8). 3 independent *Dicer1*^*-/-*^ clones have reversed transcription at the major satellite after selection for viability.

*S. pombe* and the nematode *C. elegans* have RNA-dependent RNA Polymerases (RdRP) that generate double stranded templates for Dicer. Resulting small RNAs from transposons and repeats then guide histone H3K9 methylation *(15, 38)*. Mammals lack RdRP, and we did not detect DICER1-dependent small RNAs from the major satellite transcripts in wild-type mESC (Fig. S3C), although we did detect degradation products in *Dicer1*^-/-^ cells resembling primary RNAs (priRNAs) observed in *S. pombe (39)*. DICER1-dependent siRNAs from ERVs, were limited to miRNA, while 3’tRNA fragments that match ERVs accumulated to higher levels in *Dicer1*^-/-^ cells *(40)* (Fig. S3C). The lack of DICER1-dependent siRNA is consistent with the absence of an RdRP. We also performed ChIP-seq, and observed a modest reduction in H3K9me3 at some ERVs, namely IAP and ETn, as well as at LINE1 transposable element loci (Figs. S4A, S4B, S4D, and S5B), as described previously *(41)*, which might be related to the loss of microRNA. In contrast, we found a modest increase in H3K9me3 at the major satellite (Fig. S4C), which was confirmed by immunofluorescence (Figs. S5C and S5D) as well as increased HP1*β*binding (Fig. S5E). We found little change in CENPA at the minor satellite (Fig. S5F), but we did detect a 2-fold decrease of H3K27me3 at the major satellite and a reduction of the H3K27 methyltransferase EZH2 at these loci (Figs. S5G and S5H). While it is possible that the increase in H3K9me3 at the pericentromeres was guided by priRNAs *(39)*, we conclude that loss of small RNA and H3K9me3 could not account for the strong phenotypes observed in *Dicer1*^-/-^ mESCs. We therefore performed a genetic screen to determine the underlying mechanism.

### *Brd4* and *Elp3* Mutations Suppress the *Dicer1*^-/-^ Phenotype

We performed a CRISPR-Cas9 screen for genetic modifiers that rescued the *Dicer1*^*-/-*^ proliferation and viability defects. In order to avoid aneuploids and second-site suppressors in established clonal cell lines, we introduced a domain-focused single guide RNA (sgRNA) library into wild-type cells and then induced homozygous *Dicer1*^-/-^ deletion with OHT treatment (Fig. S6A). The library targeted 176 genes responsible for chromatin modification, and sequencing at d12, d16, and d20 revealed guides that were either enriched (genetic suppressors) or depleted (genetic enhancers) after selection (Table S2). Of the suppressors, the bromodomain transcription factor *Brd4* and the histone acetyltransferase *Elp3* were outliers as the strongest hits at all three timepoints (Fig. 2A). Additionally, mutations in the H3K9 methyltransferases *Ehmt2 (G9a), Ehmt1 (GLP), Suv39h1*, and *Suv39h2* were ranked as genetic suppressors while the H3K27 methyltransferase *Ezh2* was a strong enhancer (Figs. S6B). These results were consistent with increased H3K9me3 and reduced H3K27me3 at the pericentromeric satellite in *Dicer1*^-/-^ cells, but as *Brd4* and *Elp3* were clearly the strongest suppressors they were investigated further.

**Figure 2.**
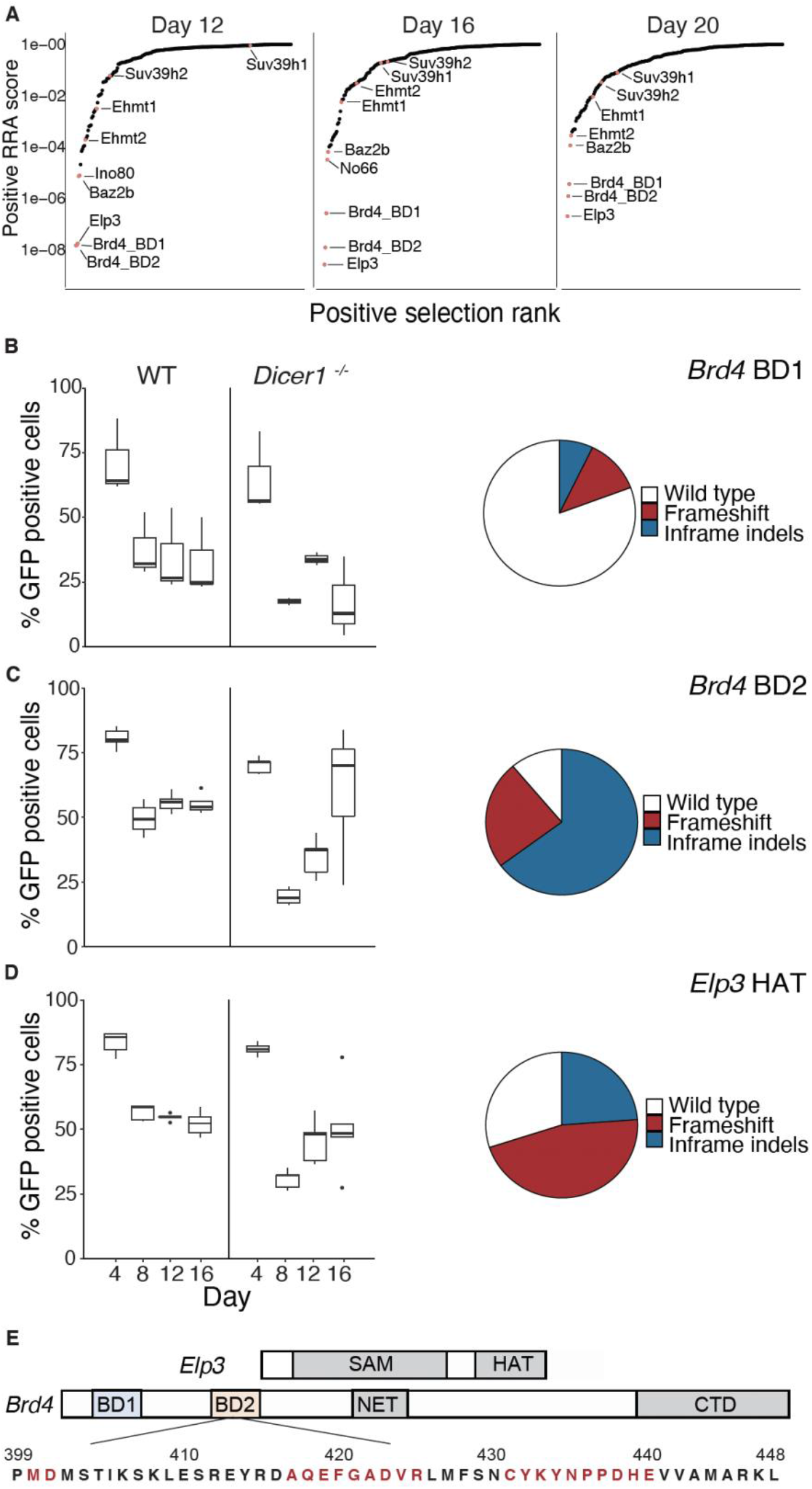
CRISPR/Cas9 suppressor screen in *Dicer1*^-/-^ cells rescues viability by in-frame mutations in *Brd4* and heterozygous frameshifts in the HAT domain *Elp3*. (A) A library of small guide RNAs (sgRNAs) targeting chromatin regulators was introduced into *Dicer1*^-/-^ cells and sequenced at three timepoints after tamoxifen-induced mutation of *Dicer1* at day 0 (d0). Individual target genes (labeled) were ranked according to positive RRA score, which ranks genes based on the positive selection of all sgRNAs targeting them. (B, C, and D) Individual sgRNAs targeting *Brd4* and *Elp3* were introduced with GFP reporter genes into *Dicer1*^-/-^ cells, and GFP positive cells were counted at the same three timepoints after tamoxifen-induced deletion of *Dicer1* (upper panels). The inferred proportion of sgRNAs targeting the first (BD1) (B) and second (BD2) (C) bromodomains of *Brd4* and the histone acetyltransferase domain of *Elp3* (HAT) (D) are shown as medians and s.d.. The relative proportion of in-frame small deletions and frameshift mutations recovered in each case are plotted as pie charts. *Brd4* BD2 mutations were strongly selected during growth, and were largely in frame. (D) The domain structure of ELP3 and BRD4 is shown as well as the amino acid sequence of the second bromodomain of *Brd4*. Amino acids missing in the small in-frame deletion alleles of *Brd4* BD2 generated by CRISPR mutagenesis are highlighted (red).

BRD4 was first described as a protein that remains bound to mitotic chromosomes *(42)*, but has since been found to globally regulate Pol II transcription at enhancer and promoter elements *(43, 44)*. BRD4 contains tandem bromodomains (BDs) that recognize acetylated lysines as well as C-terminal domains that activate transcription (Fig. 2E). However, in HeLa cells BRD4 is recruited to pericentromeric heterochromatin under heat stress *(45)*, while in *S. pombe*, the *Brd4* homolog Bdf2 regulates the boundaries of pericentromeric heterochromatin *(46)*. ELP3 is a histone acetyl-transferase (HAT) and the catalytic subunit of the Elongator complex, which also promotes Pol II transcription through chromatin *(47)* by acetylating histone H3 and H4 both in the budding yeast *Saccharomyces cerevisiae (48, 49)* and in human cell lines *(47)*. ELP3 is more thoroughly characterized as a tRNA acetyl transferase, which utilizes a radical S-adenosyl-L-methionine (SAM) domain, to modify U_34_ of some tRNAs *(50)* (Fig. 2E).

We validated these observations by introducing *Brd4* and *Elp3* individual sgRNAs in parallel together with a green fluorescent protein (GFP) reporter gene. We observed enrichment of GFP positive cells following induction of *Dicer1* perturbation consistent with mutations in these genes being suppressors (Figs. 2B, 2C, and 2D). We found that targeting the second bromodomain (BD2) of *Brd4* was more effective than targeting the first (BD1). At the conclusion of the *Dicer1*^*-/-*^ timecourse, the majority of BD1 alleles in GFP positive cells yielded a wild type amino acid sequence (Fig. 2B) suggesting a negative selection of deleterious alleles, while the majority of BD2 alleles were small in-frame deletions (Fig. 2C). This was consistent with the essential function of BD1 *(51)* and suggested that viable BD2 alleles rescued growth. The amino acids frequently deleted in BD2 are highly conserved and important in BRD4 function (Fig. 2E). In contrast, frameshifts were predominant in *Elp3* suggesting heterozygous loss of the HAT domain rescued growth, while maintaining the essential N-terminal SAM domain (Fig. 2D). We examined RNA and protein levels of BRD4 and ELP3 in *Dicer1*^-/-^ mESCs and observed little or no change (Figs. S6C and S6D) suggesting neither was a target of microRNAs. We then generated clonal single and double mutant lines using the same sgRNAs. We recovered *Brd4* mutants with heteroallelic indels in BD2 and *Elp3* mutants with heterozygous frameshift mutations in the HAT domain. These lines suppressed the viability defects of *Dicer1*^-/-^ mESCs in a luciferase-based metabolism (MT) viability assay (Figs. S6E and S6F), performing better than induced cells due to clonal selection (Figs. 3A and 3B). Additionally, the defects were rescued by *Brd4* siRNAs (Fig. 3E), though siRNAs targeting *Elp3* did not suppress the proliferation defect (Fig. 3E), presumably because they also reduced the function of the essential SAM domain. Strikingly, inhibiting Pol II elongation with low concentrations of *α*-amanitin also rescued viability (Fig. 3C), while this inhibition had no effect on wild type cells (Fig. S7C). Most significantly, the *Dicer1*^-/-^ viability defect was strongly rescued by the small molecule inhibitor JQ1, which specifically inhibits BRD4 and its paralogs (Fig. 3D).

**Figure 3.**
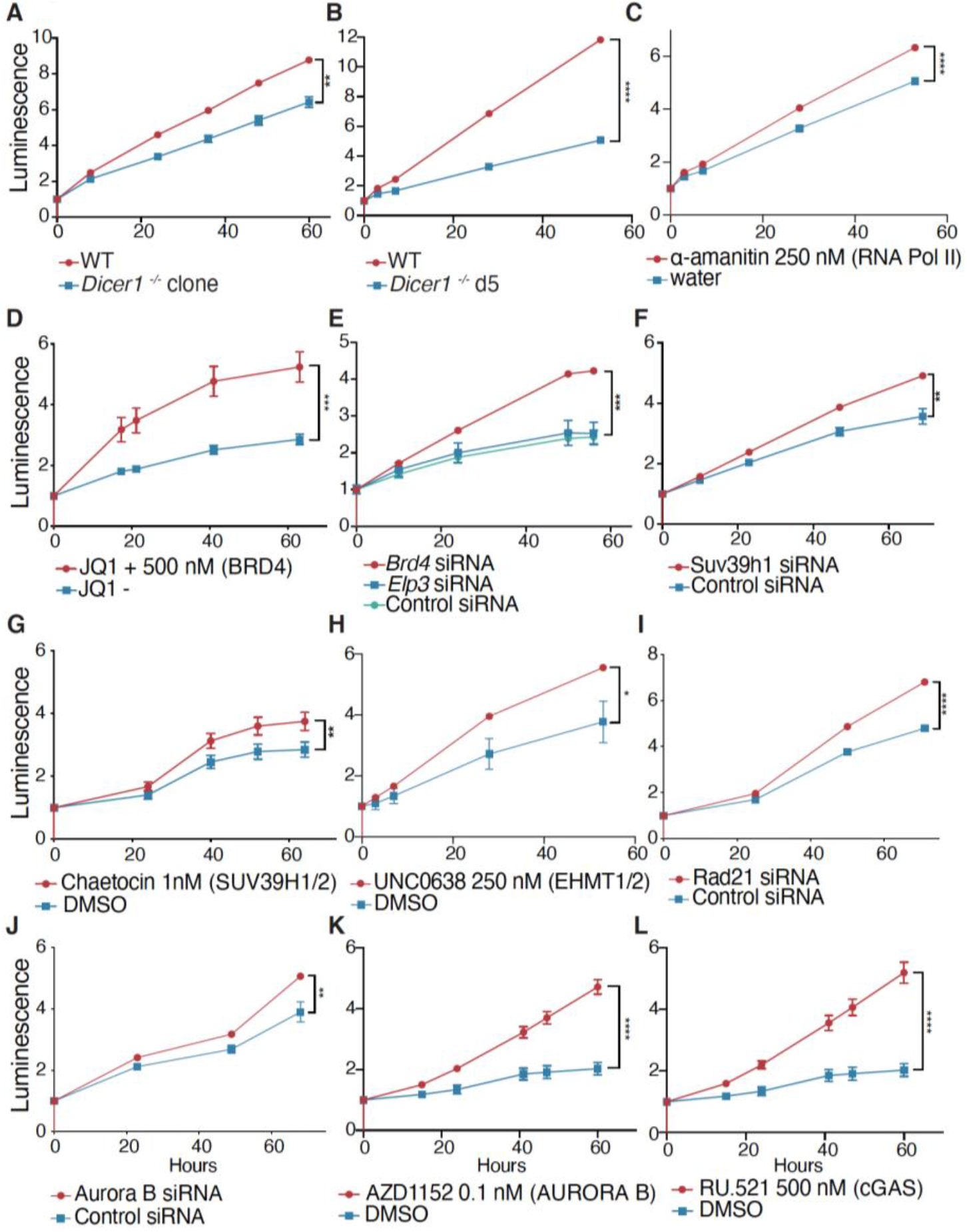
Rescue of *Dicer1*^-/-^ mESCs with siRNA and with small molecules targeting suppressors (A-B) MT proliferation and viability assay of uninduced wild type, compared with either *Dicer1*^-/-^ clones (A) or *Dicer1*^-/-^ induced cells over days 4-7 (B). *Dicer1*^-/-^ mESCs have a strong viability defect. (C-L) MT assays reveal that the proliferation and viability defect of *Dicer1*^-/-^ induced cells over days 4-7 was rescued by targeting RNA Pol II with *α*-amanitin (C), BRD4 with JQ1 (D), *Brd4* or *Elp3* with siRNAs (E), *Suv39h1* with siRNAs (F), SUV39h1/2 with the inhibitor chaetocin (G), EHMT1/2 with the inhibitor UNC0638 (H), *Rad21* with siRNAs (I), *Aurora B* with siRNAs (J), AURORA B with the inhibitor AZD1152 (K), or cGAS with the inhibitor RU.521 (L). Luminescence is plotted over time (hours) after addition of pre-luminescent metabolite at t0. DMSO was used for small molecule delivery and slightly inhibited growth of untreated controls. One representative experiment is plotted in each panel, standard error bars are shown, but may be smaller than points, **** - p-value < 0.0001, *** - p-value between 0.0001 and 0.001, ** - p-value between 0.001 and 0.01, * - p-value between 0.01 and 0.05, t test of the final timepoints of replicate experiments.

### *Dicer1*^-/-^ Chromosomal Defects Depend on Major Satellite Transcription

In order to investigate the mechanism of suppression, we performed BRD4 ChIP-seq in wild type and *Dicer1*^-/-^ mESCs. We found a two-fold reduction of BRD4 occupancy at the vast majority of genes (Fig. S8A) making them unlikely targets for rescue by *Brd4* loss-of-function mutations. Intriguingly, one feature with much higher levels of BRD4 was a 38kb portion of the major satellite repeat found in the mm10 reference genome sequence at the end of chromosome 9 (Fig. 4A), which matched 20 of the top 40 significantly differential BRD4 peaks in *Dicer1*^-/-^ clones (Table S3). Next, we determined whether transcript levels were significantly altered upon *Brd4*^*BD2-/-*^ or *Elp3*^*HAT+/-*^ mutation in *Dicer1*^-/-^ clones. *Brd4*^*BD2-/-*^ or *Elp3*^*HAT+/-*^ mutations alone had only mild effects on transcription (100-200 mostly down-regulated genes), but shared a significant number of targets (28), including known BRD4 targets *Lefty1* and *Lefty2 (52)*. Further, the mutation of *Brd4* or *Elp3* in combination with *Dicer1* resulted in differential expression of many transcripts relative to *Dicer1* single mutants (Fig. S8D), more than half of which were shared. Intersection of the datasets revealed that more than half of the differentially expressed genes had a BRD4 peak immediately upstream (<10 kb) (Fig. S8E). 97 genes were upregulated upon *Dicer1* mutation, downregulated upon *Brd4* and *Elp3* mutation, and were located near a BRD4 ChIP-seq peak, suggesting they were direct targets of both DICER1 and BRD4.

**Figure 4.**
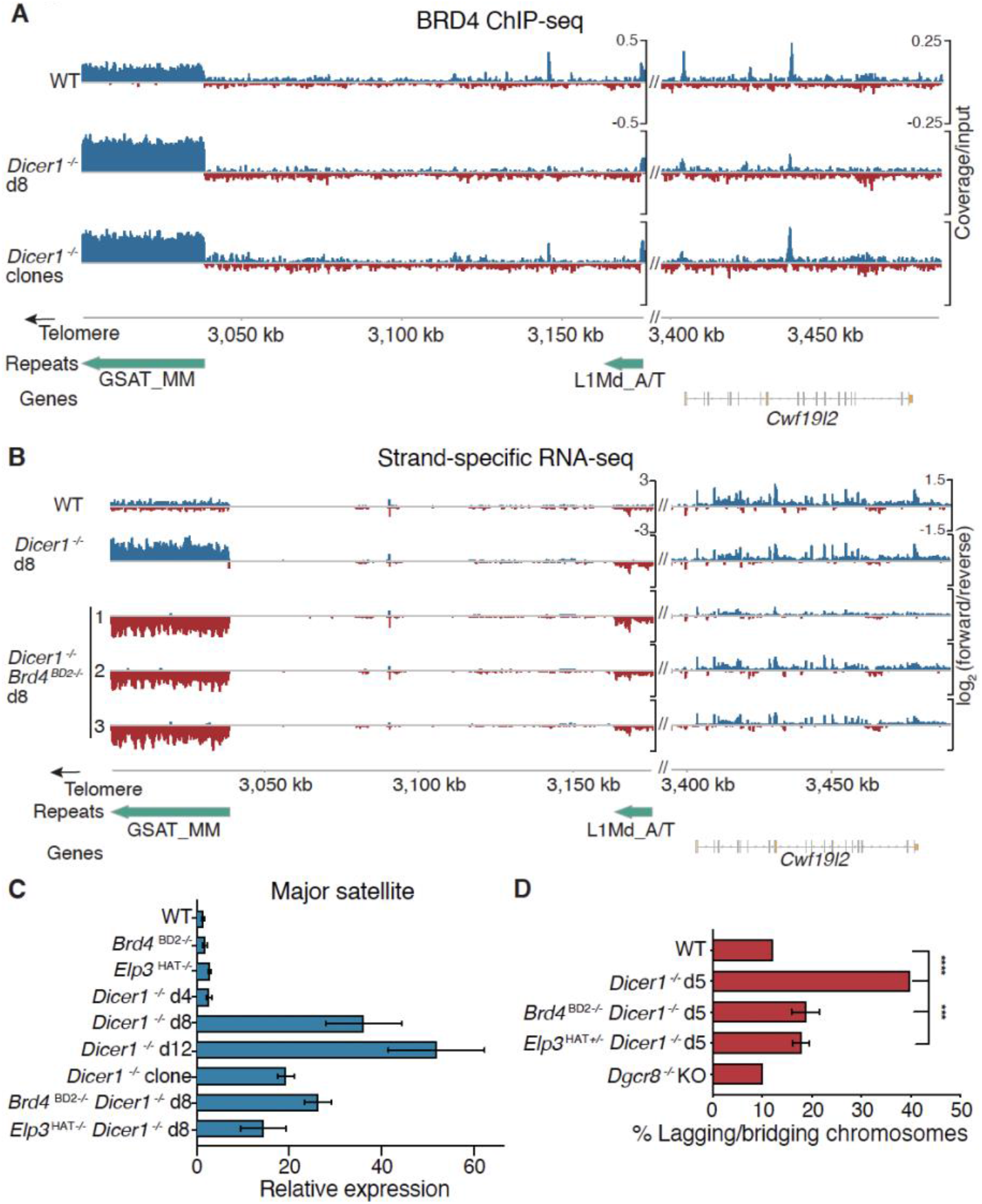
BRD4 drives expression of satellite sequences in *Dicer1*^-/-^ mESCs and suppressors reduce chromosome defects (A) BRD4 ChIP-seq coverage normalized to input plotted over the same major satellite repeat region at the end of chromosome 9 as in Fig. 1. BRD4 is enriched over the satellite repeat 8 days after deletion of *Dicer1 (Dicer1*^-/-^ d8) and in viable *Dicer1*^-/-^ clones. (B) Strand-specific transcriptional upregulation of the major satellite repeat (GSAT_MM) on chr9 is reversed in *Dicer1*^*-/-*^ *Brd4*^*BD2-/-*^ cells. The log_2_ normalized ratio of forward (blue) and reverse (red) strand is plotted. One representative replicate out of 3 is plotted for WT and *Dicer1*^*-/-*^ cells 8 days after *Dicer*1 deletion (d8). 3 independent *Dicer1*^*-/-*^ cultures have reversed transcription at the major satellite after *Brd4* mutation in the BD2 domain, 8 days after *Dicer*1 deletion. The WT and *Dicer1*^-/-^ d8 cultures are the same replicates from Figure 1. (C) RT-qPCR confirms a strong accumulation in *Dicer1*^-/-^ mESCs of transcripts from the major satellite, which is reduced in *Brd4*^*BD2-/-*^ or *Elp3*^*HAT+/-*^ doubles or *Dicer1*^-/-^ clones. Fold change was calculated relative to *Actb* and uninduced wild type controls. Error bars are standard error. (D) Chromosomal defects are reduced in *Dicer1*^-/-^ mESCs with *Brd4*^*BD2-/-*^ or *Elp3*^*HAT+/-*^ mutations. The WT, *Dicer1*^*-/-*^ d5, and *Dgcr8*^*-/-*^ data is reproduced for comparison with Fig. 1B (n > 100, **** - p-value < 0.0001, *** - p-value between 0.0001 and 0.001, Fisher’s exact test).

One of these direct targets that stood out was the major satellite repeat transcript. The abundance of the major satellite transcripts increased dramatically over the culture time-course, but was reduced in viable clones and in *Dicer1*^-/-^ d8 cells with *Brd4* or *Elp3* mutations (Fig. 4C). This transcript was also strongly downregulated in transcriptomes from *Brd4*^*BD2-/-*^ *Dicer1*^-/-^ and *Elp3*^*HAT +/-*^ *Dicer1*^*-/-*^ clonal double mutants (Fig. S8B). Strikingly, the mutation of *Brd4* reversed strand-specific transcription of the satellite transcripts in cultured *Dicer1*^*-/-*^ cells, closely resembling transcription in viable clones (Fig. 4B). Along with reduced BRD4 occupancy at satellite loci (Fig. S9A), and a reduction in elongating Pol II (Fig. S9B), chromosomal defects of *Dicer1*^*-/-*^ cells were also rescued by *Brd4*^*BD2-/-*^ and *Elp3*^*HAT +/-*^ (Fig. 4D). We further detected a substantial reduction of RAD21 at both the major and minor satellite loci in *Brd4*^*BD2-/-*^ *Dicer1*^-/-^ double mutants (Fig. S9C) consistent with recent findings that BRD4 interacts directly with RAD21 in human cells *(53)* and in *D. melanogaster (54)* as well as with NIPBL in humans *(53, 55)*. *RAD21* encodes cohesin, while NIPBL encodes the cohesin loading complex, which is responsible for proper chromosome cohesion and subsequent segregation at mitosis.

### Dicer-dependent Genome Stability via BRD4 is Deeply Conserved

In quiescent Dicer mutant *S. pombe* cells, viability is partially restored by mutation of the H3K9 methyltransferase Clr4 and its interacting partners, as well as the HP1 homolog Swi6, and by viable alleles of the spindle assembly checkpoint Ndc80, required on G_0_ entry *(18)*. In mESCs, we found that mutations in the H3K9me3 methyltransferases *Suv39h1/2, Ehmt1(GLP)*, and *Ehmt2(G9a)*, were also weak suppressors of *Dicer1*^-/-^ viability defects (Fig. 2A-C). Targeting *Suv39h1* with siRNA (Fig. 3F) and sgRNA (Fig. S7K) significantly alleviated the *Dicer1*^-/-^ viability defect, as did the *Suv39h1* inhibitor chaetocin *(56)* (Fig. 3G) and the *Ehmt1/2* inhibitor UNC0638 *(57)* (Fig. 3H). The majority of H3K9me2/3 is found at the pericentromere, and is thought to recruit the chromosome passenger complex, comprising AURORA B kinase, SHUGOSHIN 1, SMC1, SMC3, RAD21, and NIPBL *(33)*. AURORA B kinase phosphorylates members of the NDC80 complex *(58)* and centromeric cohesin *(59, 60)*, and both inhibition *(61)*, and overexpression of AURORA B *(62)*. RAD21 is a core component of the cohesin complex that confers sister chromatid attachment during mitosis and is maintained specifically at the centromere until anaphase *(63)*. Targeting either of these genes with siRNAs ameliorated the proliferation defects of *Dicer1*^-/-^ cells (Figs. 3I and 3J), as did targeting AURORA B with the pharmacological inhibitor AZD1152 *(64)* (Fig. 3K), with little effect in wild type cells (Fig. S7). We found that all three classes of suppressors—transcription, H3K9 methylation, and chromosome segregation— significantly reduced the incidence of lagging and bridging chromosomes at anaphase in *Dicer1*^-/-^ mutant clones (Fig. 4D) or in siRNA knockdowns in *Dicer1*^-/-^ cultured cells (Fig. S9D). In fact, we found that inhibiting the cytoplasmic dsDNA sensor cGAS with the small molecule RU.521 *(65)* essentially rescued the viability and proliferation defects of *Dicer1*^-/-^ mESCs (Fig. 3L) at concentrations that reduced viability of wild type mESCs (Fig. S7J). Thus, the reduction in cell viability observed in *Dicer1*^-/-^ mESCs is due in large part to the generation of cytosolic dsDNA caused by lagging chromosomes in mitosis.

In *S. pombe*, homologs of *Brd4* are encoded by Bdf1 and Bdf2, though Bdf2 more closely resembles *Brd4*. Bdf1 and Bdf2 are genetically redundant in *S. pombe (66)* and the *bdf1*Δ*bdf2*Δ double-mutant is inviable (Fig. S10A), as are *Brd4* knockout mutants in mammalian cells. Perhaps for this reason, mutants in Bdf1 and Bdf2^BRD4^ have not been recovered in fission yeast genetic screens for Dicer suppressors *(18)*. We constructed a collection of Bdf2 mutant strains (Fig. S10B) to inactivate specifically BD1 or BD2. Strikingly, only *bdf2*^BD1Δ^ displayed synthetic lethality with *bdf1*Δ (Fig. S10A), showing that, as in mammalian cells, the essential function of BET proteins is centered on BD1 *(67)*. We generated double deletion mutants of Dicer with *bdf2*Δ. The double mutant strains suppressed sensitivity to the microtubule poison thiabendazole (TBZ) (Fig. 5A), the accumulation of pericentromeric transcripts (Fig. 5C), and lagging chromosomes (Fig. 5D). Furthermore, because chromosome segregation defects cause a loss of viability on G_0_ entry, we found higher viability of the *dcr1*Δ*bdf2*Δdouble mutants (Figs. 5B and S11A). We constructed a yeast mutant equivalent to our *Brd4* CRISPR alleles in mammalian cells —a small in-frame deletion of 7 amino-acids in BD2 (Fig. 5E)—which was fully viable in the *bdf1Δ* background (Figs. S10A and S10B). Strikingly, this *dcr1*Δ *bdf2*^CR-BD2^ double-mutant was comparable to *dcr1*Δ *bdf2*Δ in TBZ sensitivity, G_0_-entry, and pericentromeric silencing (Figs. 5A, 5B, 5C and S10C). The triple-mutant *dcr1*Δ *bdf1*Δ *bdf2*^CR-BD2^ displayed nearly complete suppression of *dcr1*Δ phenotypes, with loss of TBZ sensitivity, near-wild-type viability on G_0_-entry, and transcriptional silencing of pericentromeric repeats. As in mammalian cells, pericentromeric transcripts in Dicer mutants had strong strand-specificity, which was reversed in the triple-mutant (Fig. 5C). These results indicate that Dcr1 centromeric function is deeply conserved in mESCs and fission yeast, both in cycling cells and upon G_0_ entry, and distinguishes between bromodomains BD1 (essential function) and BD2 (silencing function) of the transcription factor BRD4.

**Figure 5.**
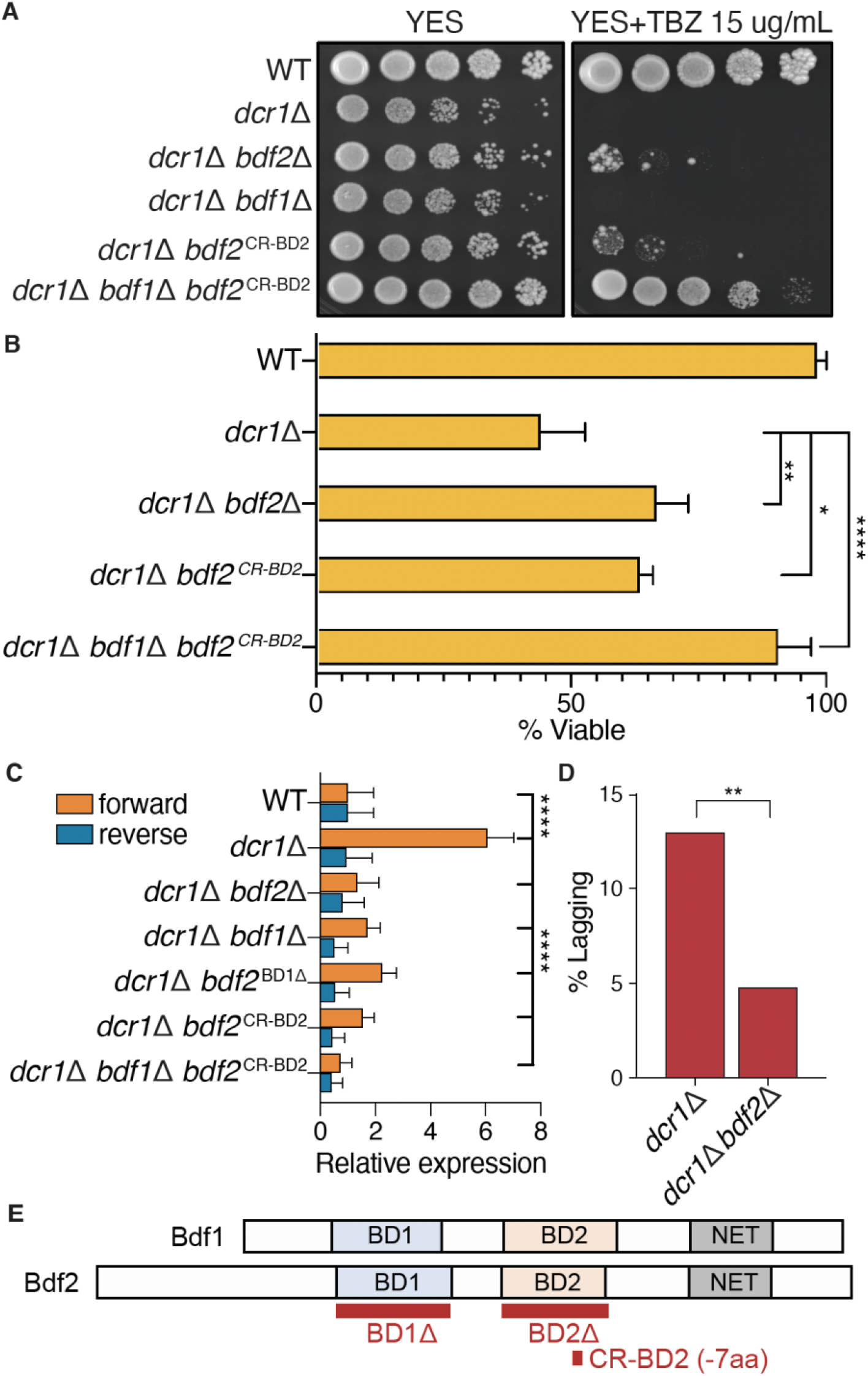
The genetic interaction of *Dicer1* with *Brd4* is conserved in *S. pombe* (A) Spot growth assays (10-fold dilutions) on media supplemented with the microtubule poison thiabendazole (TBZ). Mitotic defects of *dcr1*Δare partially suppressed by *bdf2*Δand *bdf2*^CR-BD2^, restoring viability, and fully suppressed in the triple-mutant *dcr1*Δ *bdf1*Δ *bdf2*^CRBD2^, which mimics the 7 amino acid deletion in BD2 found in *Dicer1*^*-/-*^ ES cells (Fig. 2E). (B) The loss of viability at G_0_-entry seen in dcr1Δ mutants is also suppressed in *dcr1*Δ *bdf2*Δ And *dcr1*Δ *bdf2*^CRBD2^, and is fully rescued in the triple-mutant *dcr1*Δ *bdf1*Δ *bdf2*^CRBD2^. (C) Forward strand *dh* pericentromeric transcripts accumulate to high levels in *dcr1Δ* mutants relative to the reverse strand, but are strongly reduced in double mutants with *bdf1Δ* and *bdf2Δ*, reversing strand preference. Silencing was fully restored in *dcr1*Δ *bdf1*Δ *bdf2*^CRBD2^. **** - p-value < 0.0001 – t-test. (D) Knockout of *Bdf2* in *bdf2Δ* reduces the lagging chromosome phenotype of *dcr1*Δ mutants. ** - p-value between 0.001 and 0.01 – Fisher’s exact test. (E) Schematic representation of BD1 and BD2 deletions used.

## DISCUSSION

In fission yeast, Dicer acts on RNA substrates to release Pol II from the pericentromere *(14, 15, 68)* and from other sites of collision between the transcription and replication machineries *(12, 13)*. In ES cells, we have shown that the *Dicer1*^-/-^ viability defect is due to the recruitment of BRD4 to the centromeric satellite repeats, and can be rescued by hypomorphic mutations in *Brd4* and *Elp3*. BRD4 is thought to resolve transcription replication collisions *(69)* and we found that the *Dicer1*^-/-^ viability defect can also be partially rescued by inhibiting Pol II. In support of our findings, nuclear-localized DICER1 has been found associated with satellite repeats and their transcripts, and with the nuclear protein WDHD1 in complex with Pol II *(29)*. In *Drosophila melanogaster* DCR2 physically associates with the carboxy terminal domain (CTD) of Pol II and this interaction promotes pericentromeric heterochromatin formation *(70)*. In human HEK293 cells, DICER1 physically associates with Pol II in a dsRNA-dependent fashion and prevents the accumulation of endogenous dsRNAs from satellite repeats *(71)*.

Suppression of the *Dicer1*^-/-^ phenotype strongly implies transcription as the underlying mechanism. We found strand-specific accumulation of major satellite transcripts in *Dicer1*^-/-^ mESCs that was reversed by *Brd4*^*BD2-/-*^ mutations and by selection for viability that generated our clonal lines. Similar reversals have been observed on transition between G_1_ and S/G_2_ phases of the cell cycle in mouse embryos up to the 4-cell stage, and so transcript reversal likely reflects the exit from G_1_ arrest in *Dicer1* ^-/-^ ES cell clones *(72)*. Transcription of the reverse strand may be toxic because of collision with DNA replication, which may occur predominantly in one direction as the repeats themselves have inefficient origins of replication *(12)*. Intriguingly, BRD4 interacts with PCNA, which is highly enriched on the lagging strand *(73)*. This could account for strand specificity of BRD4-dependent transcripts, which DICER1 would normally remove from the lagging strand to avoid collisions during replication. In the absence of RdRP, DICER1 may recognize a non-canonical substrate to release Pol II in ES cells. Consistent with the lack of small RNAs, there was no decrease in H3K9me2/3 at the pericentromere, as there was at retrotransposons that match small RNAs *(41)*.

We used our suppressors to further dissect the mechanism underlying the *Dicer1*^-/-^ phenotype. We found a direct relationship between transcription (BRD4) and cohesion (RAD21/AURORA B) that suggests transcriptional suppressors may act through the cohesin complex, as in *S. pombe (74, 75)*. Ectopic expression of satellite transcripts in mouse and human cells has been found to cause chromosome mis-segregation and genome instability in tumors *(76)*. The mechanism is thought to involve interaction of major satellite transcripts with the MCM complex required for replication, and replication stalls in the S phase of the cell cycle *(76)*, very much like the *Dicer1*^-/-^ mESCs analyzed here. These observations suggest a re-evaluation of the high frequency of *Dicer1* mutants in cancer *(77–79)*, in which chromosomal abnormalities lead to increased genome instability. We find that chromosomal phenotypes activate cytoplasmic DNA sensors, such as cGAS/STING, which respond to the presence of micronuclei *(80)* and generate a type I interferon response that reduces proliferation and leads to apoptosis *(81)*. Our data indicates this mechanism is the cause of the proliferation and viability defects of *Dicer1*^-/-^ mESCs.

Aberrant accumulation of major satellite RNA has also been shown to recruit heterochromatin factors SUV39H1 *(82–84)* and HP1 *(85)* to the pericentromere. However, the mutation of *Brd4*, which reduces transcript levels significantly, did not result in loss of H3K9me3. Instead, H3K9me3 accumulation in *Dicer1*^-/-^ mESCs more closely resembles the phenotype of yeast *dcr1*Δquiescent cells in which rDNA accumulates H3K9me2 and relevant small RNAs are not detected *(18)*. The arrest of *Dicer1*^-/-^ ES cells in G_1_ suggests that they may enter a non-dividing G_0_-like state that affects viability. Major satellite transcripts accumulate primarily in late G_1_ *(72, 86)* and G_1_ arrest is a prerequisite for entry into G_0_ in fission yeast and mammalian cells *(79, 87, 88)*. If the loss of DICER1 does force the cells into a suspended G1 or G0-like state and DICER1 is required to maintain this state as it is in *S. pombe*, then second site mutations or epimutations would be required to exit this state and resume growth.

### A Conserved Mechanism for Dicer and Transcription at the Centromere

We have identified suppressors of the *Dicer1*^-/-^ ES cell proliferation, viability and chromosomal defects in transcriptional activators *(Brd4* and *Elp3)*, H3K9 methyltransferases, and chromosome segregation factors *(Aurora B* and *Rad21)*. These same classes of suppressors rescued viability defects in Dicer deletion mutants in *S. pombe (18)*. The most significant upregulated non-coding RNA in *Dicer1*^-/-^ mESCs was the major satellite RNA, directly comparable to pericentromeric transcripts in *S. pombe*. Thus, the mechanism by which Dicer regulates transcription, even in the absence of RdRP and small RNAs, is conserved from fission yeast to mammals. The novel role we identified for BRD4 in pericentromeric transcription was also conserved in *S. pombe*. Strikingly, this conservation held true for specific bromodomains, with Bdf2-BD1 being essential in the absence of Bdf1, while Bdf2-BD2 is involved in suppression of *dcr1*Δ. Mutants in the centromeric H3K14 acetyltransferase Mst2 suppress *dcr1*Δ in a similar fashion *(89)*, and H3K14A mutants lose pericentromeric silencing *(90)*. In *S. pombe, bdf2*Δ mutants alleviate DNA damage accumulation at the S-phase checkpoint, and suppress hydroxyurea sensitivity in checkpoint mutants *(66)*. *dcr1*Δ mutants accumulate stalled RNA pol II and DNA damage at S phase *(12)*, and suppression by *bdf2*Δ results in reduction of reverse-strand pericentromeric transcription, lower amounts of stalled RNA pol II and reduced DNA damage. This allows maintenance of heterochromatin through the cell cycle and ensures its mitotic inheritance, as in *mst2*Δ *(89)*. In cancer cells, BD1 is also essential, while BD2 has specific roles in inflammation, autoimmune disease, and specific cancer subtypes. Moreover, BD2 specific inhibitors are promising therapeutic agents for these conditions with minimal side effects *(91)*. BD1 has higher affinity for acetylated lysines on histone H4, while BD2 has a higher affinity for H3 acetyl lysines *(92)*, consistent with a key role for H3 modifications in centromeric silencing in *S. pombe*. These conserved genetic and mechanistic interactions with transcription, DNA replication, histone modification, and sister chromatid cohesion likely contribute to “Dicer syndrome”, in which Dicer mutations pre-dispose cancer and viral infection *(79)*. A potential therapeutic role for BRD4 and BD2 inhibitors is suggested by these interactions.

## ACKNOWLEDGMENTS

We thank Edith Heard (EMBL, Heidelberg, Germany) and Iku Okamoto (Kyoto University) for the inducible *Dicer1* mutant lines, and Chris Vakoc (Cold Spring Harbor Laboratory) for providing advice and reagents and for reading the manuscript. M.J.G. was supported by a Bristol-Myers Squibb Fellowship from the Cold Spring Harbor Laboratory School of Biological Sciences. A.J.S. was supported by NIH grant R01GM076396 to R.A.M. This work was supported by the Howard Hughes Medical Institute and the Gordon and Betty Moore Foundation (GMBF3033). The authors acknowledge assistance from the Cold Spring Harbor Laboratory Shared Resources, which are funded in part by the Cancer Center Support Grant (5PP30CA045508).

## AUTHOR CONTRIBUTIONS

Conceptualization, M.J.G., B.R., A.J.S., and R.M.; Methodology, M.J.G., K.C., and B.R., Formal Analysis, M.J.G., B.R., and J.I.S.; Investigation, M.J.G., B.R., J.I.S., A.L., and A.S.J. Writing – Original Draft, M.J.G.; Writing – Review & Editing, M.J.G., B.R., and R.M.; Funding Acquisition, R.M.

### DATA AND MATERIALS AVAILABILITY

All data, materials, and methods are available upon request.

## METHODS

### EXPERIMENTAL MODEL AND SUBJECT DETAILS

#### Cell Lines and Tissue Culture

Mouse embryonic stem cells, including *Dicer1* ^*flx/flx*^ lines, were grown in 2i conditions. The 2i medium consists of: 250 mL Neurobasal medium (Gibco 21103-049), 250 mL DMEM/F12 (Gibco 11320-033), 2.5 mL N2 supplement (Gibco 17502-048), 5 mL B27 supplement (Gibco 17504-044), 5 mL GlutaMAX (Gibco 35050-061) and 3.3 mL 7.5% BSA Fraction V (Gibco 15260-037). Just prior to adding the media to the cells, the following additional supplements were added: PD0325901 to 1 *μ*M (Stemgent 04-0006), CHIR99021 to 3 *μ*M (Stemgent 04-0004), 2-Mercaptoethanol to 100 *μ*M (Gibco 21985-023), and Mouse LIF to 10^3^ units/mL (EMD Millipore ESG1106). Culture dishes were coated with gelatin (1X Attachment Factor – ThermoFisher S006100) for a minimum 15 minutes, gelatin was removed, and dishes were allowed to dry before cells were plated. Cells were split with TrypLE Express Enzyme (Gibco 12605-010) for 5 minutes at 37 degrees Celsius. Cells were collected in enough media for a 10-fold dilution of the TrypLE Express and pipetted up and down a minimum of 10 times to achieve a single cell suspension. Cells were then spun for 5 minutes at 150 x g and resuspended in media for counting and plating. Cells were frozen in freezing media: 10% DMSO (Sigma D2650), 50% ES-FBS (Gibco 16141-002), and 40% cells in media. Inducible *Dicer1* knockout cells were treated with hydroxytamoxifen (OHT – Sigma H7904) at a final concentration of 1 *μ*M. For immunofluorescence, cells were grown on poly-L-lysine coated coverslips in 6 well dishes (Corning 354085).

Lenti-X 293T cells were grown in DMEM with 10% FBS and pen/strep. All cell lines used in this study were grown at 37 degrees Celsius and 5% CO_2_. All mouse embryonic stem cells used are male.

## METHOD DETAILS

### Transfections

In the case of siRNA transfection, the Lipofectamine RNAiMAX was used (ThermoFisher 13778030) according to the manufacturer’s protocol. The siRNAs were the OnTarget Pools targeting genes of interest from Dharmacon. In the case of Cas9/sgRNA transfection, Lipofectamine CRISPRMAX (ThermoFisher CMAX00001), TrueCut Cas9 (ThermoFisher A36497), and TrueGuide sgRNAs (ThermoFisher custom) were used. The Lipofectamine CRISPRMAX protocol was followed for transfection preparation. In all cases, transfection reagents were prepared ahead of time and added to wells. Cells were then split and added to the transfection-containing media (reverse transfection).

### DNA/RNA Isolation and cDNA Preparation

Cells were collected by splitting as described and pelleted. To isolate genomic DNA, the Purelink Genomic DNA Mini Kit (K182001) was used as directed. Genomic DNA was eluted in TE and stored at −20 degrees Celsius. Concentration was measured with the Nanodrop. RNA isolation was performed on either fresh pellets or fresh pellets were resuspended in the appropriate amount of TRIzol (ThermoFisher 15596018), snap frozen, and stored at −80 degrees Celsius until the protocol could be finished. Cells were resuspended in the appropriate amount of TRIzol and the manufacturer’s protocol was followed as directed, save for the use of 80% ethanol for the washing steps to increase retention of small RNAs. RNA was resuspended in ultra-pure water and stored at −80 degree Celsius. RNA concentration was measured by Nanodrop and quality was measured with the Bioanalyzer. For cDNA preparation 1 microgram aliquot of pure RNA was DNase treated and reverse transcribed using the Superscript IV Vilo ezDNAse kit (ThermoFisher 11766050).

### RT-qPCR and ChIP-qPCR

In general qPCR was performed with Taqman probes. For qPCR on genomic DNA, genomic DNA was quantified using Nanodrop or Qubit and diluted to 5 nanograms per microliter. The Taqman CNV Assay and *Tfrc* CNV Assay control were run together in triplicate in 10 microliter reactions on a 384 well plate with the 2X Genotyping Mastermix (ThermoFisher 4371355) and analyzed with the ΔΔCt method. For qPCR on cDNA, the cDNA was generally diluted at least 1 to 20 and 1 microliter of diluted cDNA was used in 10 microliter reactions on a 384 well plate in a reaction with the Taqman Assay, *Actin* or *Gapdh* control assay with a different fluorophore, and the Taqman Fast Advanced Master Mix (ThermoFisher 4444557). The run was analyzed with the ΔΔCt method. For ChIP-qPCR, ChIP DNA was diluted along with the H3 and Input controls at least 1 to 10. For each run a standard curve was run to test efficiency of the primers. The reactions were 10 microliters in 384 well plates with PowerUp SYBR Green Master Mix (ThermoFisher A25918). Each run was tested for efficiency and then a percent input calculation was used to determine enrichment.

### Western Blot

Cell pellets were washed twice with PBS and resuspended in an appropriate amount of RIPA buffer (ThermoFisher 89900) and incubated for 15 minutes on ice. Samples were centrifuged at 14,000 x g for 15 minutes and the supernatant was transferred to a new tube. Protein concentration was measured with a microplate BCA assay in triplicate (Pierce 23252). Equal amounts of protein (at least 15 micrograms) was loaded on a 4-20% gradient gel (Bio-Rad 4561096) and run until loading dye reached the bottom of the gel. The proteins were transferred onto nitrocellulose membranes using the Trans-Blot Turbo Transfer system. A Ponceau stain was then performed to verify the transfer. Gels were often divided to probe for different sized proteins. Blocking was done in 5% Milk in TBST. Primary antibodies were incubated for at least an hour at room temperature or overnight at 4 degrees Celsius. Washes were performed with TBST. Secondary antibodies were incubated for at least 1 hour at room temperature. The secondary antibody was detected with the SuperSignal West Pico Plus Chemiluminescent Substrate (ThermoFisher 34577) and imaged with the Bio-Rad ChemiDoc MP. In the need for stripping and re-probing, Restore Western Blot Stripping Buffer was used (ThermoFisher 21059).

### Cell Proliferation and Viability Assays

For cell proliferation and viability assays the Promega RealTime-Glo MT Cell Viability Assay (Promega G9711) was used. The assay was used according to the protocol, with the two reagents diluted to 1X in a white 96 well plate that had been coated with 1X Attachment Factor. Cells were then added to well and allowed to stabilize for at least 1 hour before the first reading. Luciferase readings were performed periodically over a period of 72 hours in a 96 well plate reader at 37 degrees Celsius with a 250 millisecond integration time at a height of 1 millimeter above the plate. Readings were done in at least 5 sites in the well to minimize noise. The assay was analyzed by normalizing all readings to the initial reading (relative luminescence in the figures) and each timepoint was plotted over time.

### BrdU Cell Cycle Analysis

BrdU Flow Cytometry Cell Cycle Analysis was performed with a FITC BrdU Flow Kit (BD Pharmingen 559619). Cells were cultured normally and then dosed with an appropriate amount of BrdU for at least 1 hour prior to harvest. The Flow Kit protocol was followed as directed and the Bio-Rad SE3 Cell Sorter was used.

### Immunofluorescence Staining and Quantification

Cells were grown on glass coverslips in 6 well dishes. Media was removed, cells were washed twice with PBS, and then treated with 4% paraformaldehyde in PBS for 10 minutes at room temperature. Cells were washed 3 times in PBS and treated with a quenching solution (75mM NH_4_Cl and 20 mM Glycine in PBS) for 10 minutes. Fixed coverslips then permeabilized with PBS-Triton X-100 0.1% for 2 minutes on ice. Blocking and antibody dilutions were performed with 5% BSA. The primary antibody was diluted as directed and applied for 1 hour at room temperature. The secondary antibody was diluted as direction and applied for 45 minutes at room temperature. Prolong Gold with DAPI (ThermoFisher P36931) was used for mounting.

### Cloning sgRNA and Cas9 Plasmids

In order to generate a lentiviral Cas9 construct, we cloned a PGK promoter and mCherry fluorescent reporter into the Lenti-Cas9-puro vector (gift from Ken Chang/Vakoc Lab). In order to generate sgRNA lentiviral vectors we first ordered sense oligos with a CACCG 5’ overhang and antisense oligos with a AAAC 5’ overhang and a C 3’ overhang. The oligos were phosphorylated and annealed with T4 PNK (NEB M0201S) in a thermocycler with the following conditions: 37 degrees Celsius for 30 minutes and then a ramping down from 95 degrees Celsius to 25 degrees Celsius at 5 degrees Celsius per minute. Annealed oligos were cloned into a Bsmb1 digested Lenti-sgRNA-GFP-LRG plasmid (gift from Ken Chang/Vakoc Lab) with T4 DNA ligase (NEB M0202S) for 30 minutes at room temperature. A 1:200 dilution of annealed oligo was used (1 uL) with 25 nanograms of digested plasmid (1 uL) in a 10 microliter ligation reaction. Lentiviral plasmids were transformed into Stbl3 (ThermoFisher C737303).

### Lentiviral Production and Infection

The PSPAX2, VSVG, and transfer plasmids were prepared according to standard bacterial culture practices in Stbl3 (ThermoFisher C737303). General bacterial culture practices were followed except the cultures were grown at 30 degrees Celsius overnight to reduce recombination. Lentiviruses were made with Lenti-X 293T cells (Takara 632180) cultured in DMEM with 10% FBS and pen/strep. Prior to transfection, cells were plated on a 15 centimeter dish coated with gelatin as described. Cells were grown until 90% confluence. Transfection of PSPAX2, VSVG, and transfer plasmids was performed with Lipofectamine 3000 according to the forward transfection protocol. Lentiviral containing media was collected at two timepoints depending on the yellowing of the media. The lentivirus was then isolated from the spent media using the Lenti-X Concentrator (631231) and following the protocol as directed. Pelleted virus was resuspended in 2i media, aliquoted, snap-frozen on dry ice, and stored at −80 degrees Celsius. Infections in ES cells were performed by adding the concentrated lentivirus to attached cells with 8 ug/mL of DEAE-Dextran (Sigma D9885). Lentivirus was removed after 24 hours and then the cells were cultured normally.

### CRISPR Screen

The Chromatin Modifier sgRNA library was prepared by spreading and growing the bacteria on large agar plates. The plates were scraped and a Qiagen Maxiprep (Qiagen 12162) was performed to isolate the lentiviral plasmids. Lenti-X 293T cells were transfected with the library as described and lentivirus was collected with the Lenti-X concentrator as described. Six replicate 15 centimeter (cm) dishes of ES cells were then infected with the concentrated lentiviral particles at a calculated representation of 500 events for each sgRNA. After 2 days the six dishes were split into two separate 15 cm dishes each and one of the two was treated with OHT to induce the *Dicer1* mutation. At this time, the GFP% in the population was monitored to ensure the infection was at a multiplicity that would produce on average less than 1 sgRNA per cell. The individual 15 cm dishes were grown over a period of 3 weeks. Each time the dishes were split genomic DNA was collected for sgRNA amplification. At the end of the growth period genomic DNA was isolated with the Qiagen Blood and Tissue Kit (Qiagen 69504). The sgRNA loci were amplified from the amount of genomic DNA necessary to maintain the representation of the library ∼500x using the high fidelity Q5 polymerase (NEB M0492) and Illumina-compatible barcoded primers were used in a second PCR to create libraries for each replicate at each timepoint. Libraries were quantified with the KAPA Illumina Quantification Kit (KAPA KK4824).

### Single sgRNA Infection and Flow Cytometry

Single sgRNA vectors were constructed and lentiviral particles were generated as described. The Cas9-expressing *Dicer1* ^*flx/flx*^ cell line was infected as described before. Two days after infection, the cells were split. At this time one third of the cells were plated and kept cultured as normal, one third of the cells were induced to mutate *Dicer1* with OHT for 24 hours, and one third of the cells were washed in PBS, run over a strainer cap flow cytometry tube, and analyzed for EGFP signal on the Bio-Rad SE3 flow cytometer. The cells were then cultured as one *Dicer1* wild type population and one *Dicer1* mutant population and every four days the cells were split and cells were again analyzed by flow cytometry for EGFP signal. At the end of the timecourse genomic DNA was collected as described for amplicon sequencing.

### Amplicon Sequencing Library Preparation

Genomic DNA from single sgRNA lentiviral timecourse experiments was isolated as described. Primers flanking the sgRNA-targeted region that contained Illumina sequence overhangs were used in a PCR reaction with genomic DNA and Q5 high fidelity polymerase (NEB M0492). A second PCR with primers containing barcodes and targeting the Illumina overhangs of the first PCR was performed with a minimal number of cycles to avoid amplification bias. The PCR-generated libraries were gel purified to remove primers. Libraries were quantified with the KAPA Illumina Quantification Kit (KAPA KK4824).

### Small RNA Sequencing Library Prep

Small RNAs were isolated from total RNA using Novex TBE-Urea Gels (ThermoFisher EC6885BOX). Libraries were constructed with the TruSeq Small RNA Kit (Illumina RS-200-0012). Libraries were quantified with the KAPA Illumina Quantification Kit (KAPA KK4824).

### Whole Genome Sequencing Library Preparation

Genomic DNA was isolated as before. DNA was fragmented with Covaris ultrasonication with default settings for the generation of fragments of 350 basepair average size. Libraries were constructed with the TruSeq DNA PCR-Free Kit (Illumina 20015962). Libraries were quantified with the KAPA Illumina Quantification Kit (KAPA KK4824).

### RNA Sequencing Library Preparation

RNA was isolated with TRIzol as described and DNase treated. When total RNA was sequenced, 1 microgram RNA was first depleted for ribosomal RNA using the rRNA Depletion Kit (NEB E6310). Total RNA libraries were then prepared using the NEBNext Ultra II Directional Kit (NEB E7760). Libraries were quantified with the KAPA Illumina Quantification Kit (KAPA KK4824).

### ChIP and ChIP Sequencing Library Prep

Cells were grown in 2i conditions as detailed on 15 centimeter plates. 37% formaldehyde is added to a final concentration of 1% in the media and incubated at room temperature for 10 minutes. The reaction was quenched with for 5 minutes at room temperature with the 10X glycine buffer from the SimpleChIP Kit (Cell Signaling 56383). The protocol for the SimpleChIP Kit was followed as directed. Chromatin was fractionated using the 1 mL millitubes on the Covaris with optimized custom settings to achieve an average size of 250 basepair fragments. ChIP was performed with recommended dilutions of antibodies and control H3 antibodies. An input control was also generated for each sample. ChIP-sequencing libraries were constructed with the NEBNext Ultra II DNA Library Prep Kit (NEB E7645S). Libraries were quantified with the KAPA Illumina Quantification Kit (KAPA KK4824) and checked for quality on the Agilent Bioanalyzer.

### *S. pombe* strains and crosses

*S. pombe* strains were cultured in standard conditions at a temperature of 30°C. *De novo* mutant strains and tagging were generated by multiplex PCR in which the first PCR run generates ∼200 nucleotides of homology regions and a second PCR step generated the final construct. PCR products were cleaned by Qiagen PCR Purification (Qiagen 28104) and 1 microgram of DNA was used per transformation using the Frozen-EZ kit (Zymo T2001). Crosses were performed on malt extract media (ME) at room temperature, followed by random spore analysis, except for the investigation of synthetic lethality between *bdf1*Δ and *bdf2* mutant strains where tetrad analysis was performed. In each tetrad analysis experiment, a minimum of 50 tetrads were isolated and dissected on YES plates, using a Sanger MSM400 micro-manipulator. Viable colonies were genotyped, and only tetrads where the genotype of every spore could be observed or inferred were taken into consideration for measuring the viability of each genotype; this assay showed complete synthetic lethality (no recovered viable double-mutant) of the *bdf1*Δ *bdf2*Δ and *bdf1*Δ *bdf2*^BD1Δ^ genotypes, and no viability defect in the other double-mutants.

### *S. pombe* Assays

In order to induce G_0_, cells were cultured in EMM (Edinburgh minimal medium) and then shifted to EMM-N (Edinburgh minimal medium without nitrogen). Viability was determined 24 hours after G_0_ entry by isolating ≥100 single-cells on a YES plate, using a Sanger MSM400, and measuring their ability to reform a colony, as previously (Roche et al, 2016). The presence of missegregation in G_0_ entry was determined by determining the proportion “rod-shaped” cells in the culture (24h G_0_) with a hemocytometer, counting ≥100 cells. Thiabendazole (TBZ assays) were performed on YES plates containing 15μg/ml of TBZ (Sigma). Cells were grown to log phase, counted, and spotted in equal numbers across a range of dilutions. RT-qPCR for the *dg/dh* repeats was performed by first isolating RNA from log phase cells with the Quick-RNA Fungal/Bacterial Kit (R2014). RNA was then DNase treated and reverse transcribed with the Superscript IV Vilo ezDNase Kit (ThermoFisher 11766050), using centromere-specific primers (p30-F and p30-R) for the strand-specific RT-qPCR, and random hexamers for non-specific RT-qPCR. The resulting cDNA was then used for qPCR using iQ SYBR Green mix (Bio-Rad), using p30 and act1 primers in the non-stranded reaction, and p30 in the stranded reaction.

## QUANTIFICATION AND STATISTICAL ANALYSIS

### Image Quantification

Intensity of immunofluorescence staining was quantified by first acquiring at least 5 Z-stacks with identical acquisition conditions for each sample across multiple coverslips. Fiji image analysis software *(93)* was used to calculate a maximum intensity projection and then measure the mean fluorescence intensity across all nuclei in an image. Lagging and bridging phenotypes were quantified by assaying each anaphase for either phenotype by visual inspection.

### Cell Proliferation and Viability Assays Statistical Analysis

All statistical analyses comparing treatment were performed using GraphPad Prism software. Unpaired t-tests were used to compare data consisting of two population means at the final timepoints of replicate assays.

### CRISPR Screen Sequencing Analysis

Sequences were de-multiplexed and the adapters were trimmed from both ends with Flexbar *(94)*. Mapping was performed with Bowtie *(95)* on a Bowtie index built with the sgRNA library sequences with the default settings and ‘--norc’ to prevent alignment to the reverse complement of the sgRNA sequences. MAGeCK analysis package was used to determine sgRNA enrichment/depletion as well as gene ranking in the screen *(96)*.

### Amplicon Sequencing Analysis

Reads were trimmed with Trimmomatic *(97)* to remove all excess Illumina adapter. Reads were collapsed using the FastX toolkit and analyzed for CRISPR-generated mutations with a custom script.

### Small RNA Sequencing Analysis

Raw reads were first trimmed with Cutadapt *(98)* to clip Illumina 3’ adapters, while retaining the 4nt-long degenerate sequences at the 5’ and 3’ end of each insert. Reads with an insert size between 12-60 nt were subsequently filtered using Gordon Assaf’s FASTX-Toolkit to discard low-quality reads with Phred scores less than 20 in 10% or more nucleotides. PCR duplicates were removed using PRINSEQ *(99)* (derep option 1) and degenerate sequences were extracted and appended to the name of each read using UMI-tools *(100)*. Preprocessed reads were first aligned to calibrator sequences and then either to *Mus musculus* mm10 UCSC genome using Bowtie2 *(101)*, which assigns multi-mapping reads in an unbiased way. Aligned reads were filtered for 0-2 mismatches using Samtools *(102)* and BamTools *(103)*. For ERV and GSAT (major satellite) analysis, aligned reads were intersected with RepeatMasker annotation (http://www.repeatmasker.org) using BEDtools *(104)*. Read counts were normalized to total calibrator per library. Shaded regions in each panel display minimum and maximum values of three biological replicates for each genotype. Data visualization was performed using R and all analysis was conducted using custom Perl and Bash scripts.

### Whole Genome Sequencing Analysis

Illumina reads were trimmed for adapters with Trimmomatic *(97)*. Reads were mapped with bowtie2 with default settings to the mm10 genome *(101)*. Samtools *(102)* was used to convert, sort, and index bam files. Freebayes *(105)* was used with default settings to call SNPs in sequencing data relative to the reference *Mus musculus* mm10 genome. SNPs were filtered for quality and SnpEff *(106)* was used to annotate. Filtered and annotated SNPs were also intersected with PFAM domains with BEDtools *(104)*. Normalized coverage was calculated using deepTools *(107)*.

### RNA Sequencing Analysis

Reads were trimmed for adapters and quality with Trimmomatic *(97)*. Mapping to the mm10 genome was performed with the STAR aligner *(108)* with --winAnchorMultimapNmax and -- outFilterMultimapNmax set to 100 in order to allow reads to multimap for downstream analysis. The analysis of differential expression of both genes and repetitive elements was performed with TEtranscripts *(109)* and DESeq2 *(110)*. Gene set enrichment analysis was performed with GSEA *(111, 112)*. Intersections between RNA-seq data sets and ChIP-seq datasets were performed with custom scripts. For strand-specific analysis, trimmed reads were mapped with bowtie2 with default settings and -k 5 to allow for multimapping. Normalized, strand-specific coverage was generated using deepTools *(107)*.

### ChIP Sequencing Analysis

Reads were trimmed for adapters and quality with Trimmomatic *(97)*. Bowtie2 *(101)* was used for mapping with default settings and –dovetail, -X 600, and -k 5. The resulting SAM files were converted, sorted, and indexed with SAMtools *(102)*. Normalized coverage was calculated for library size for each sample and a relative normalized coverage was calculated for IP over input using deepTools *(107)*. Peaks were called with default settings with MACS2 *(113)*. Differential peaks were called with default settings with MAnorm *(114)*. Peaks were assigned to underlying or nearby features and a functional enrichment analysis of these features was performed with ChiP-Enrich *(115)*. The intersection of ChIP peaks/assigned features with differentially expressed RNA-sequencing results was performed with custom scripts. Normalized coverage was calculated and heatmap/profile plots were made using deepTools *(107)*.

All custom commands and scripts are available upon request.

