## Supplemental material for "Dicer promotes genome stability via the bromodomain transcriptional co-activator Brd4"

### LIST OF SUPPLEMENTARY MATERIALS

1. Figure S1 - Inducible deletion of *Dicer1* in mouse embryonic stem cells.
2. Figure S2 - Additional chromosomal phenotypes in *Dicer1*$^{-/-}$ mutants.
3. Figure S3 - *Dicer1*$^{-/-}$ mESCs have an apoptosis signature and low levels of major satellite transcript degradation products.
4. Figure S4 - Histone H3 Lysine 9 methylation is largely retained in the absence of *Dicer1*.
5. Figure S5 - *Dicer1*$^{-/-}$ mESCs retain H3K9me2/3 chromatin at transposable elements and satellite repeats, but lose H3K27me3.
6. Figure S6 - Screening and validation of *Ezh2* and *Brd4*/*Elp3* as enhancers and suppressors of *Dicer1*$^{-/-}$ viability defects.
7. Figure S7 - siRNA and small molecule inhibitors do not affect proliferation of wild type mESCs.
8. Figure S8 - BRD4 binding at genes does not change upon *Dicer1* mutation and BRD4 and ELP3 co-regulate genes and major satellite transcripts.
9. Figure S9 - Classes of suppressors interact and suppress chromosomal defects of *Dicer1*$^{-/-}$ mESCs.
10. Figure S10 - *bdf1Δ* and *bdf2Δ* in *S. pombe* are synthetic lethal via bromodomain 1 (BD1), but suppress *dcr1Δ* via BD2.
11. Table S1 – Differentially expressed transcripts in *Dicer1* mutant clones compared to uninduced wild type control.
12. Table S2 – Results of a CRISPR-Cas9 genetic screen of chromatin modifiers in *Dicer1*$^{-/-}$ mESCs.
13. Table S3 – Differential BRD4 ChIP-seq peaks in *Dicer1*$^{-/-}$ (timecourse day 8 or clones – KO) vs wild type.

Supplementary Tables S1, S2, and S3 are provided as separate excel spreadsheets.

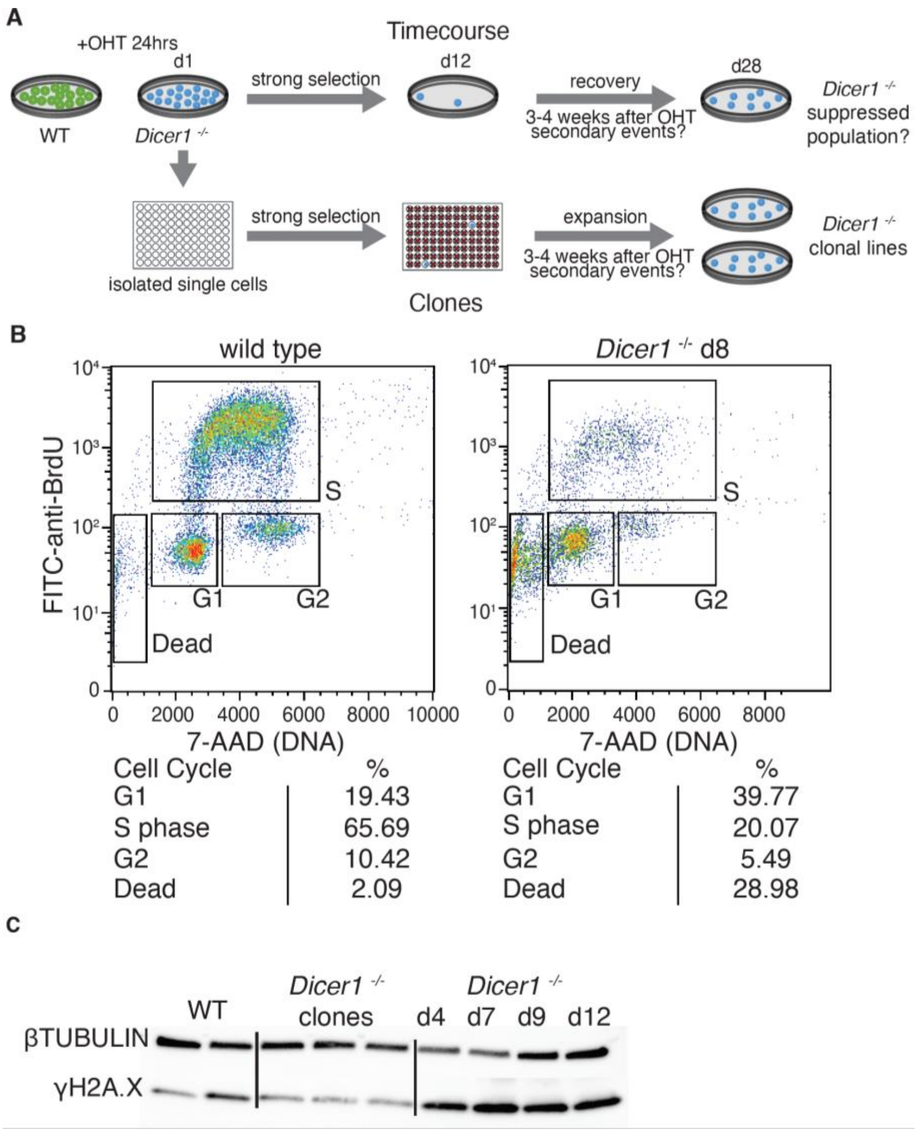


Figure S1. Inducible deletion of *Dicer1* in mouse embryonic stem cells.

(A) *Dicer1*$^{-/-}$ timecourse samples were derived from *Dicer1 flx/flx* mESCs that express a tamoxifen-inducible Cre recombinase by treatment with hydroxytamoxifen at day 0 (d0). Induction generated a population of *Dicer1*$^{-/-}$ cells that had lost both RNase III domains. These cells were grown for a period of 3-4 weeks in 2i media and resulted in stalled proliferation and apoptosis peaking at days 8-12 (d8-12) after induction. *Dicer1*$^{-/-}$ clones were derived from cells isolated immediately after induction of *Dicer1* mutation (d1). After 3-4 weeks in culture in 96 well plates, a small fraction of clones achieved a proliferation rate high enough to grow as a clonal cell line. After 4 weeks, timecourse populations were dominated by rare wild-type cells without rearrangement that were strongly selected for viability.

(B) Flow-cytometry analysis of the cell cycle in wild type and *Dicer1*$^{-/-}$ ES cells on day 8 after tamoxifen-induced deletion (*Dicer1*$^{-/-}$ d8). Cells were sorted into G1, S and G2 phase by BrdU incorporation and DNA content.

(C) DNA damage was assessed by Western blot using antibodies against $\gamma$H2A.X in the *Dicer1*$^{-/-}$ timecourse and clones.

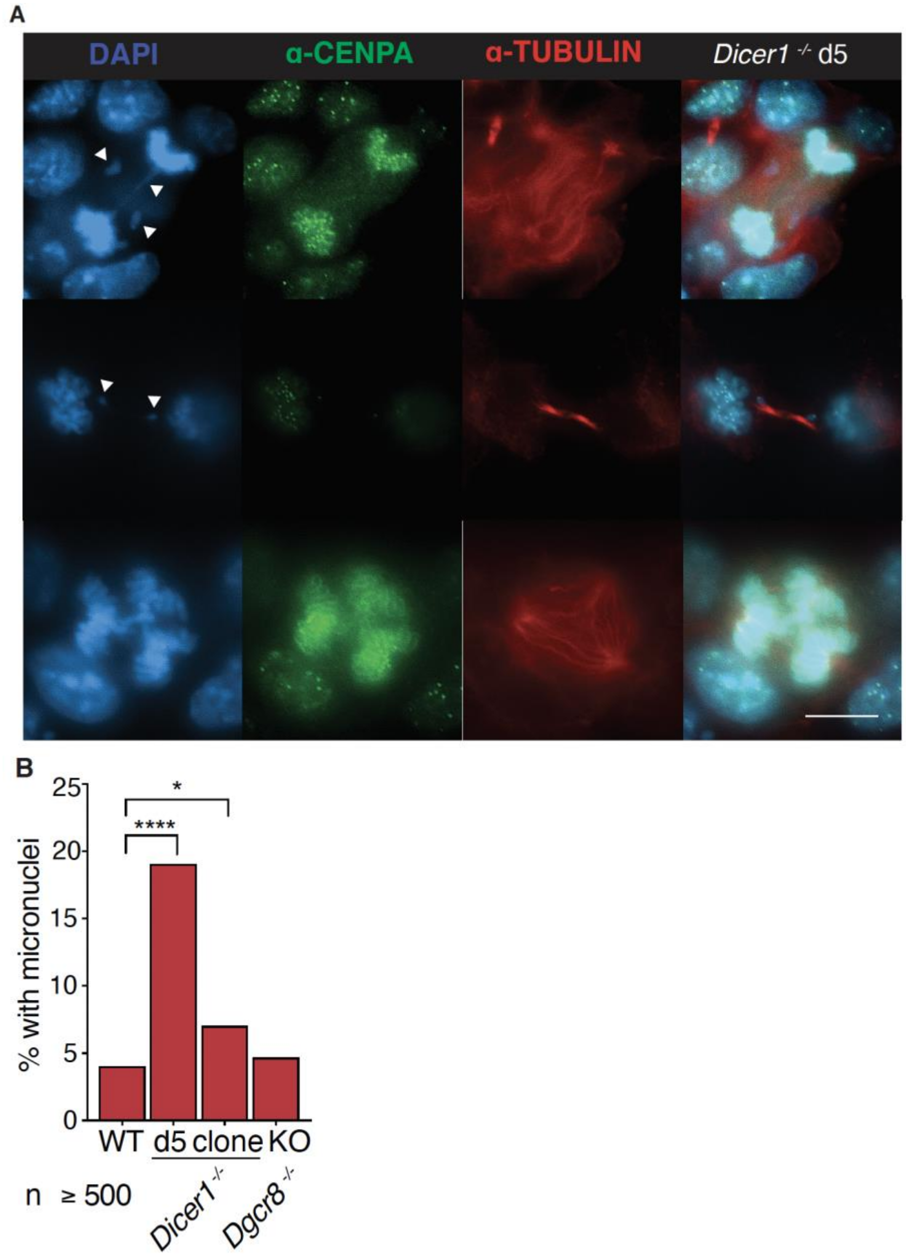


Figure S2. Additional chromosomal phenotypes in *Dicer1*$^{-/-}$ mutants.

(A) Immunofluorescence of CENPA (green) and tubulin (red) with DAPI staining (blue) shows CENPA incorporation in lagging chromosomes (top) and micronuclei (middle) and shows the underlying cytoskeletal structure of the tripolar mitoses observed (bottom). (Scale bar = 10 microns).

(B) Quantification of micronuclei at interphase (**** - p< 0.0001, * - 0.01 >p> 0.05 - Fisher's exact test). Scale bar = 10 microns (n > 500).

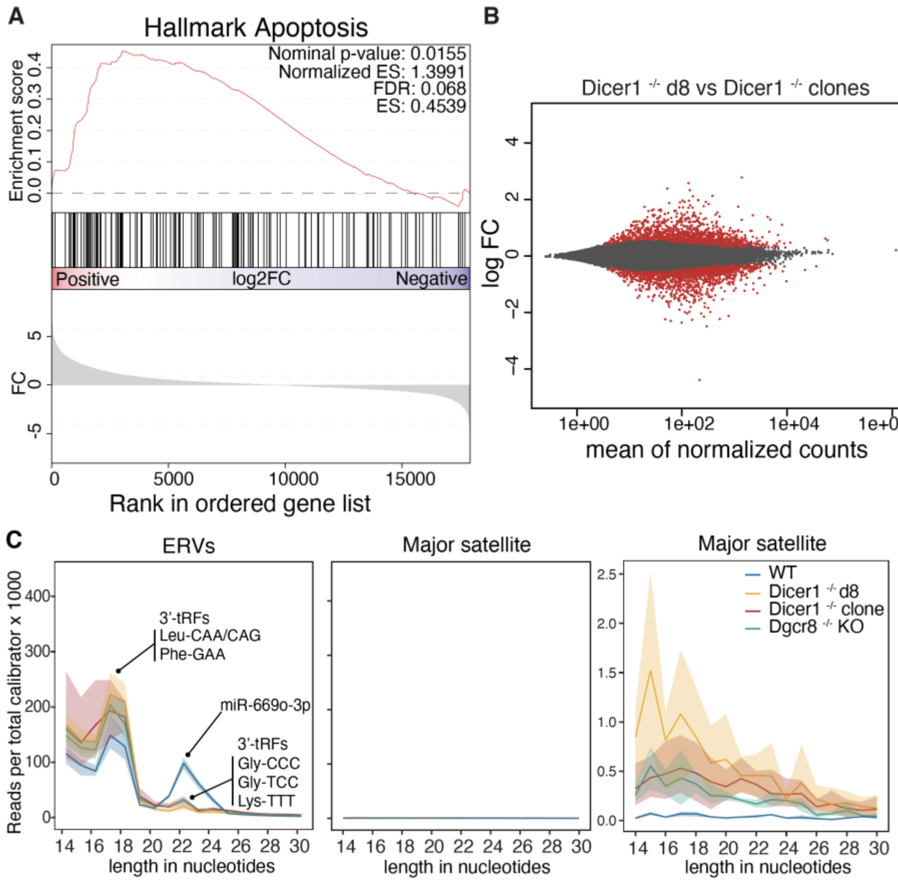


Figure S3. *Dicer1*$^{-/-}$ mESCs have an apoptosis signature and low levels of major satellite transcript degradation products.

(A) Hallmark Gene Set Enrichment analysis (GSEA) demonstrates an upregulation of apoptosis promoting factors in *Dicer1*$^{-/-}$ clones relative to wild type.

(B) Differential expression analysis identifies significant difference in transcripts altered in *Dicer1*$^{-/-}$ cells 8 days after deletion (d8) compared to mutant clones that have undergone selection for viability.

(C) Small RNA-seq identifies low levels of degradation products from the major satellite transcripts in *Dicer1*$^{-/-}$ clones. Results are plotted as counts normalized to calibrator spike-in abundance to account for the loss of microRNAs and shading indicates the range of replicate values. The right panel is a zoomed in plot of the center panel with an appropriate scale.

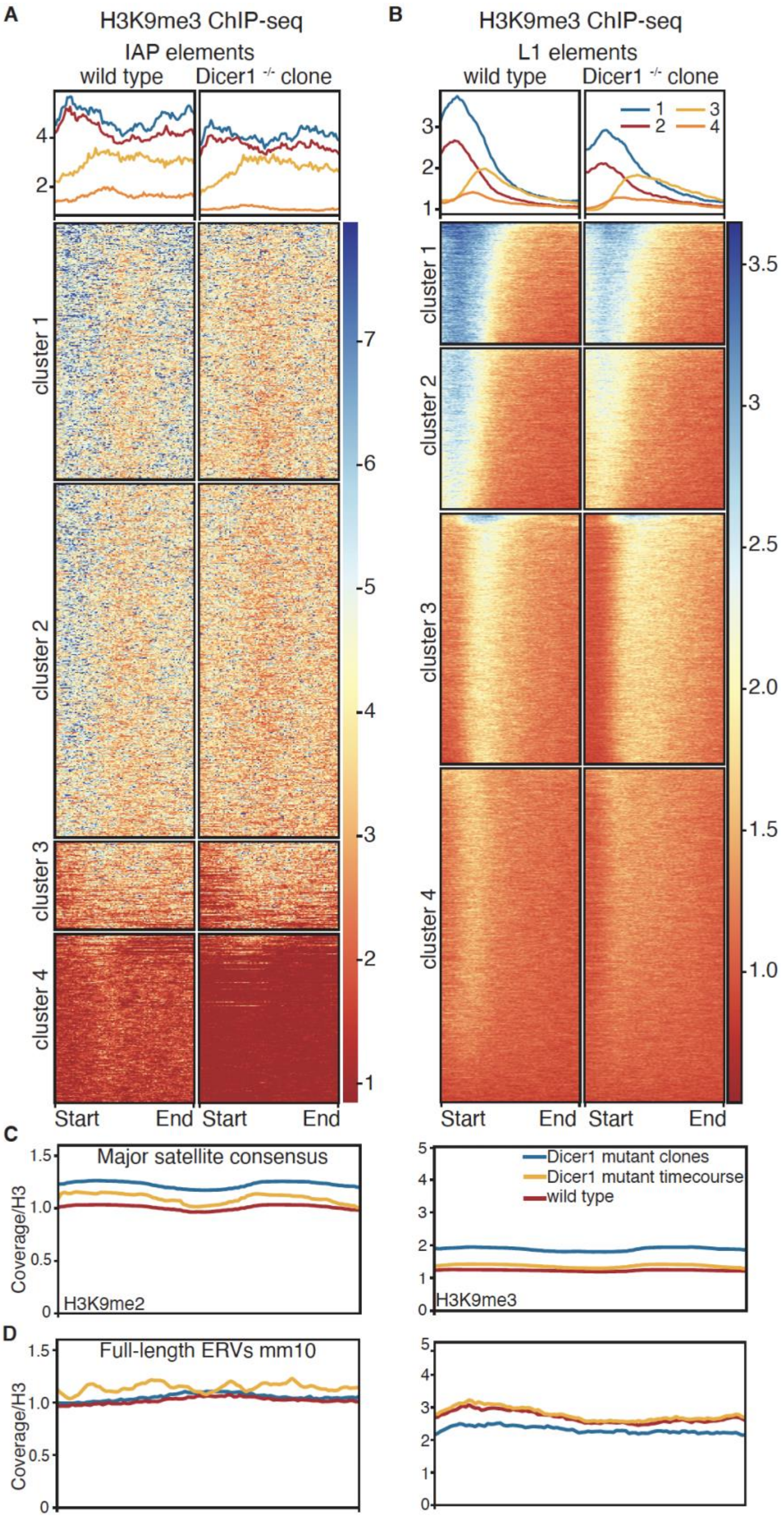


Figure S4. Histone H3 Lysine 9 methylation is largely retained in the absence of *Dicer1*.

(A and B) Clustered heatmaps of H3K9me3 normalized to input and histone H3 over full length IAP (A) and LINE1 (B) transposable elements. Some subclasses of LINE1 elements have reduced H3K9me3 in *Dicer1*$^{-/-}$ day 8 cells, but most TEs are unaffected.

(C and D) ChIP-seq reads from H3K9me2 and H3K9me3 normalized to input and histone H3 in wild type, *Dicer1*$^{-/-}$ day 8, and *Dicer1*$^{-/-}$ clones were aligned to a 2x consensus sequence of the major satellite (C) and full length ERVs (D). H3K9me2 and 3 were largely retained in the absence of *Dicer1*.

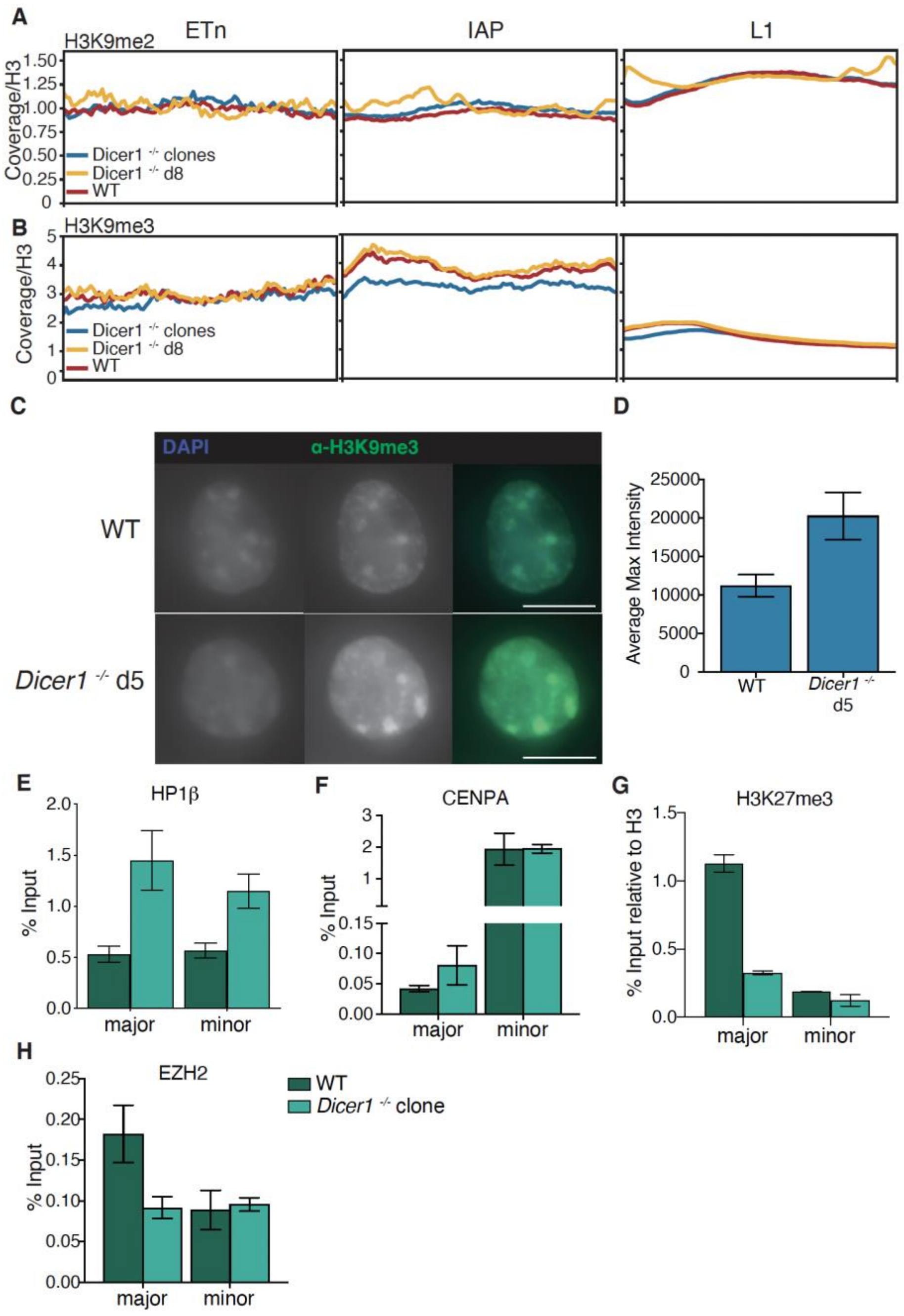


Figure S5. *Dicer1*$^{-/-}$ mESCs retain H3K9me2/3 chromatin at transposable elements and satellite repeats, but lose H3K27me3.

(A and B) The ChIP-seq coverage of H3K9me2 (A) and H3K9me3 (B) normalized to input and H3 control are plotted for wild type, *Dicer1*$^{-/-}$ timecourse day 8, and *Dicer1*$^{-/-}$ clonal line samples across all full length ETn elements (left), full length IAP elements (center), and L1 elements in mm10.

(C and D) Immunofluorescence of H3K9me3 in uninduced wild type (C) and *Dicer1*$^{-/-}$ timecourse day 5 cells (D) showed a global increase in H3K9me3 in *Dicer1*$^{-/-}$ concentrated at chromocenters (scale bar = 10 $\mu$M). This increase was quantified as the average max intensity of collapsed Z-stacks acquired with the same exposure from multiple independent slides and experiments (n > 50).

(E, F, G, H) ChIP-qPCR reveals the gain of HP1$\beta$ in *Dicer1*$^{-/-}$ clonal lines at the major and minor satellite repeat loci (E), the maintenance of centromeric identity as measured by CENPA incorporation (F), and the loss of H3K27me3 (G) and EZH2 binding (H). Results are plotted as percent input (error bars are standard error).

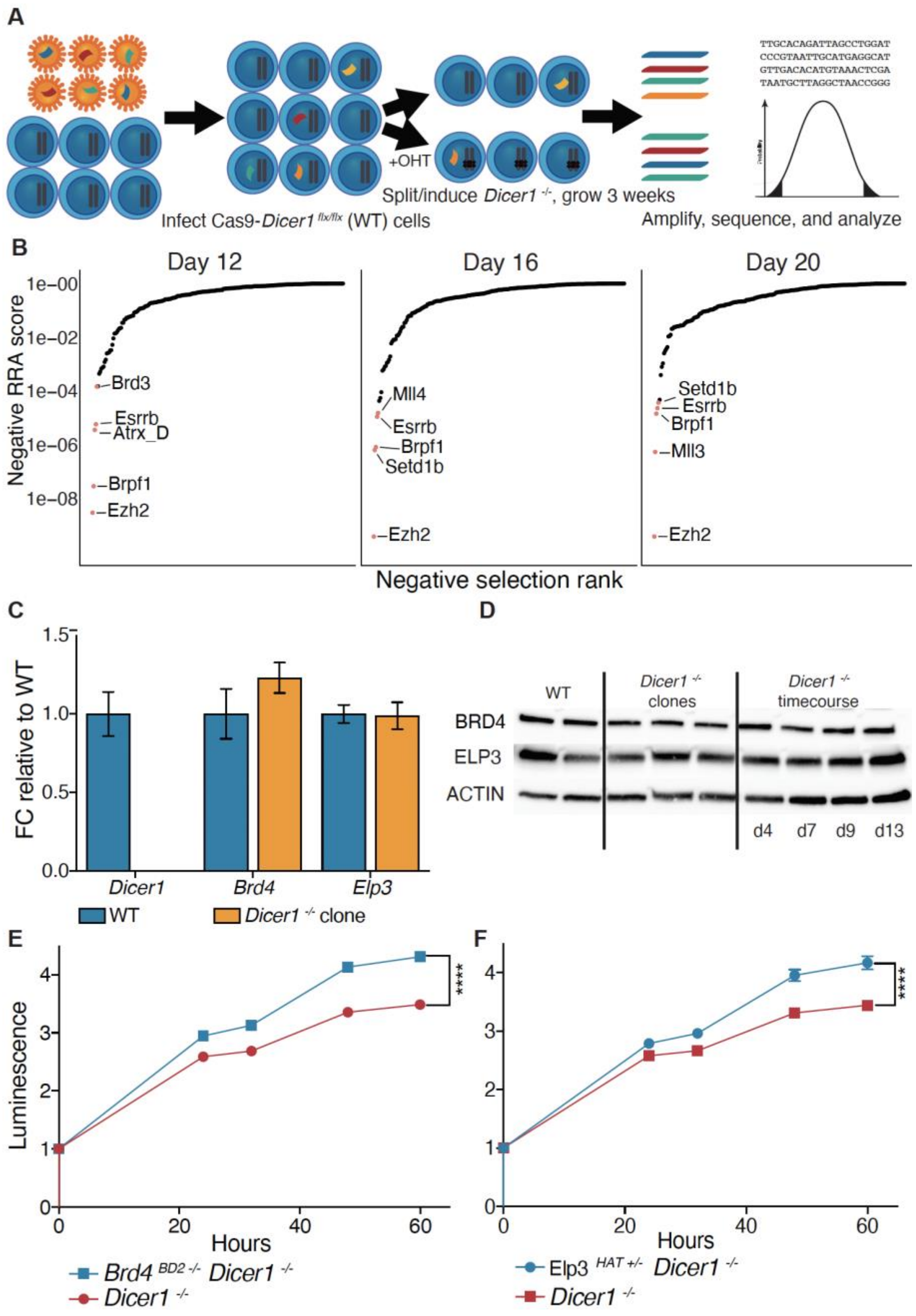


Figure S6. Screening and validation of *Ezh2* and *Brd4*/*Elp3* as enhancers and suppressors of *Dicer1*$^{-/-}$ viability defects.

(A) A CRISPR-Cas9 genetic screen was performed by transducing a *Dicer1 flx/flx* cell line that was constitutively expressing Cas9 with a domain-focused library of sgRNAs targeting chromatin modifiers in six replicate experiments. The transduced cells were split into two populations – 1. *Dicer1 flx/flx* untreated wild type replicates and 2. *Dicer1*$^{-/-}$ replicates that were treated with hydroxytamoxifen to induce recombination at the RNase III domains of both *Dicer1* alleles. The independent replicate populations were grown for a period of three weeks and genomic DNA was collected every 4 days when the populations were split. The integrated sgRNA sequences were amplified from the population and subjected to next generation sequencing to determine the relative abundances in each replicate.

(B) The results of the CRISPR-Cas9 genetic screen are plotted according to the negative RRA score, which ranks genes based on the performance of all sgRNAs targeting them, and the ranking of genes as being negatively selected. The results at the timepoints following the phenotypic selection of *Dicer1* mutation are shown.

(C and D) The relative amounts of *Brd4* or *Elp3* do not change in *Dicer1*$^{-/-}$ day 8 at either the RNA level by RT-qPCR fold change calculated with normalization to *Actb* and uninduced wild type samples (C) (error bars are SE) or the protein level as determined by Western blot (D). Therefore, they are very unlikely to be targets of miRNA.

(E and F) An MT-like proliferation and viability assay during days 4-7 of the *Dicer1*$^{-/-}$ timecourse shows a suppression of the proliferation defect of *Dicer1* by 3 independent clonal mutations in the second bromodomain of *Brd4* (E) or the histone acetyltransferase domains of *Elp3* (F). One representative experiment is plotted, standard error bars may be smaller than points, **** - p-value < 0.0001, t test of final timepoints of replicate experiments.

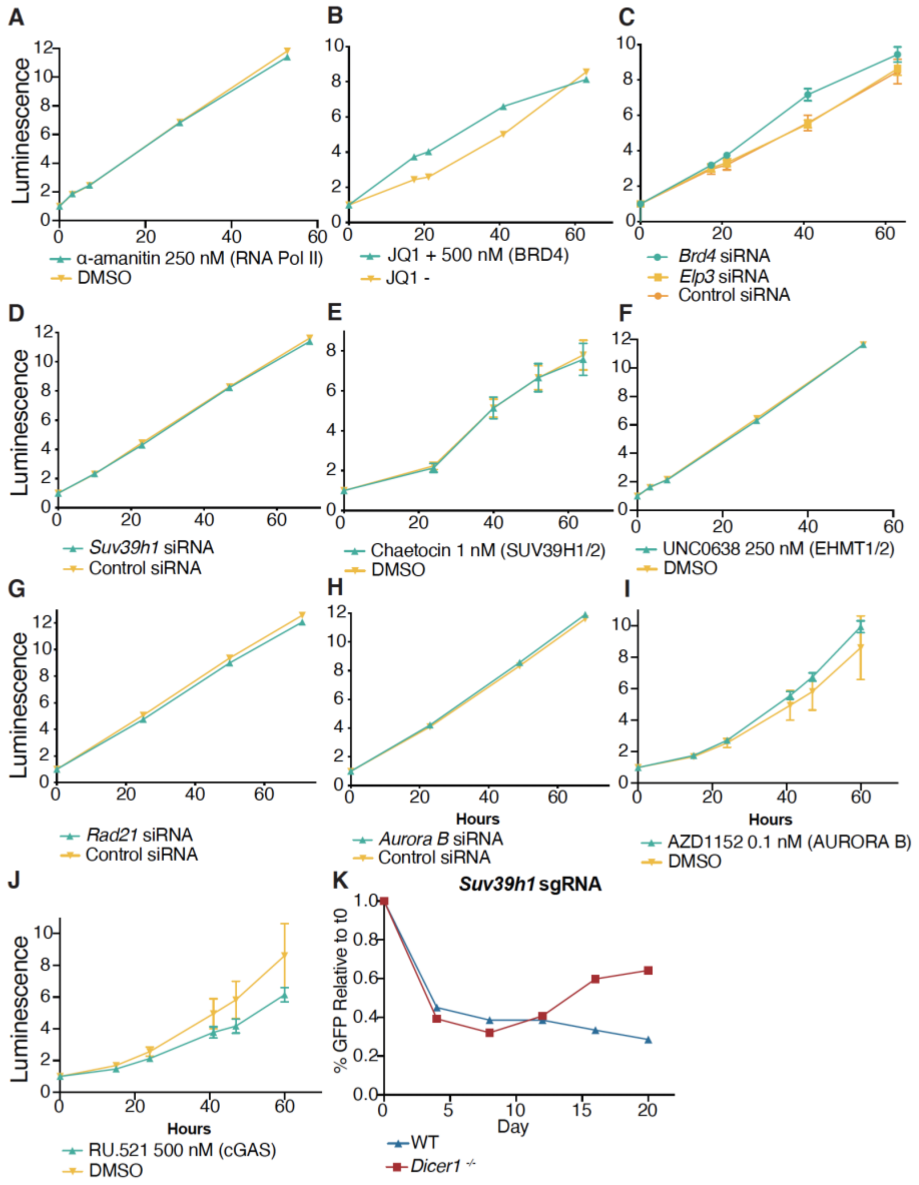


Figure S7. siRNA and small molecule inhibitors do not affect proliferation of wild type mESCs.

(A-J) An MT-like proliferation and viability assay shows siRNA and small molecule inhibitor treatments have little effect on wild type mESCs by targeting RNA Pol II with $\alpha$-amanitin (A), *Brd4* with JQ1 (B), *Brd4* or *Elp3* with siRNAs (C), *Suv39h1* with siRNAs (D), *Suv39h1/2* with the inhibitor chaetocin (E), *Ehmt1/2* with the inhibitor UNC0638 (F), *Rad21* with siRNAs (G), *Aurora B* with siRNAs (H), *Aurora B* with the inhibitor AZD1152 (I), or cGAS with the inhibitor RU.521 (J). One representative experiment is plotted, standard error bars may be smaller than points.

(K) Individual sgRNAs targeting the first exon of *Suv39h1* were introduced with eGFP reporter genes into *Dicer1*$^{-/-}$ cells, and GFP positive cells were counted at the same three timepoints after tamoxifen-induced deletion of *Dicer1* (upper panels). The inferred proportion of sgRNAs are shown as medians and s.d..

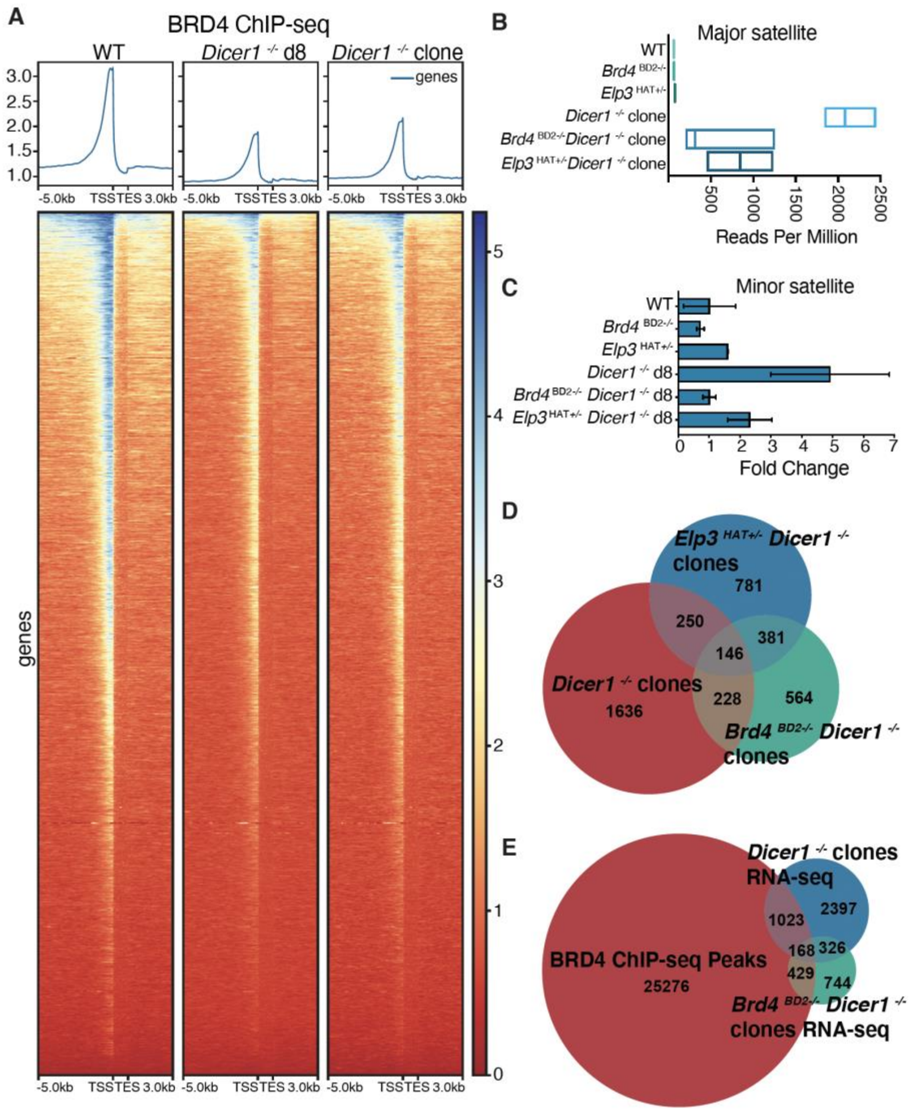


Figure S8. BRD4 binding at genes does not change upon *Dicer1* mutation and BRD4 and ELP3 co-regulate genes and major satellite transcripts.

(A) The binding profile of BRD4 at all UCSC RefSeq Genes is plotted by normalized coverage over the transcriptional units from 5 kb upstream to 3 kb downstream. This analysis showed a general reduction in binding of BRD4 to promoter regions of genes in *Dicer1*$^{-/-}$ conditions.

(B) Normalized reads per million of the major satellite transcript shows the substantial reduction in abundance in *Brd4* or *Elp3* double mutants with *Dicer1*. The line in each box represents the median.

(C) RT-qPCR confirmed an accumulation in *Dicer1*$^{-/-}$ mESCs of transcripts from the minor satellite, which was reduced in *Brd4*$^{BD2-/-}$ or *Elp3*$^{HAT+/-}$ doubles or *Dicer1* clones. Fold change was calculated relative to *Actb* and uninduced wild type controls (error bars are SE).

(D) An intersection of significantly differentially expressed transcripts upregulated in *Dicer1*$^{-/-}$ mESCs and downregulated in *Brd4*$^{BD2-/-}$ and *Elp3*$^{HAT+/-}$ double mutants demonstrate co-regulation of the majority of these transcripts by BRD4 and ELP3.

(E) An intersection of significantly differentially expressed transcripts in *Brd4*$^{BD2-/-}$ *Dicer1*$^{-/-}$ clonal lines with BRD4 ChIP-seq peaks within 10 kb of the locus identified in *Dicer1*$^{-/-}$ . Importantly, all 146 co-regulated transcripts (which included the major satellite transcript) identified in panel D were included in the 168 peaks, but only the major satellite had dramatically increased levels of BRD4 ChIP in *Dicer1*$^{-/-}$ cells (Fig. 4A).

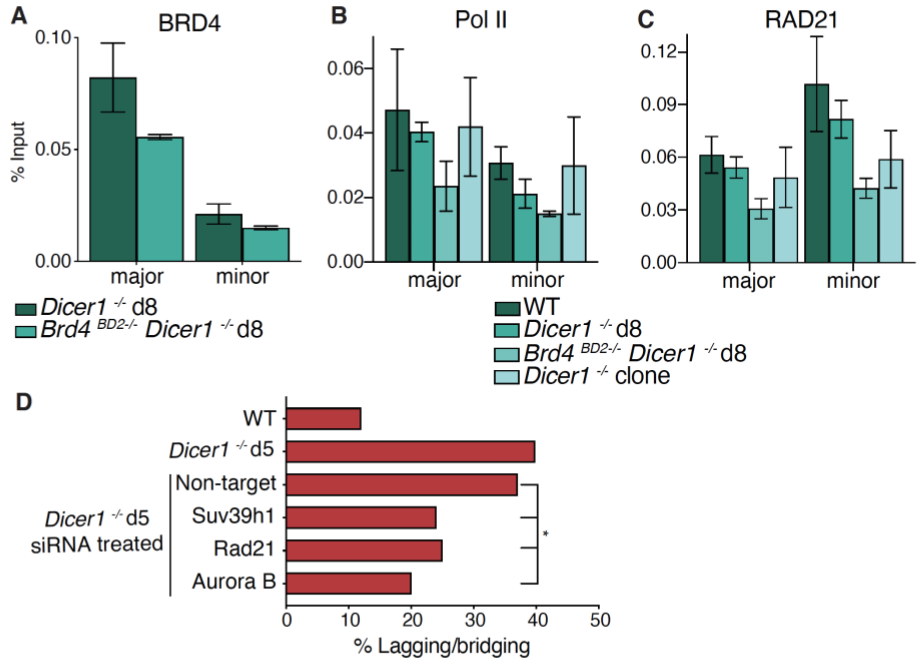


Figure S9. Classes of suppressors interact and suppress chromosomal defects of *Dicer1*$^{-/-}$ mESCs.

(A) ChIP-qPCR at the major and minor satellite repeats showed an increase in H3K9me3 in *Brd4*$^{BD2-/-}$ *Dicer1*$^{-/-}$ timecourse cells compared to *Dicer1*$^{-/-}$ timecourse cells. The results are plotted as percent input (error bars are standard error).

(B and C) ChIP-qPCR at the major and minor satellite repeats uncovered a reduction in active RNA polymerase II (Ser2/5 phosphorylated) (B) and RAD21 (C) in *Brd4*$^{BD2-/-}$ *Dicer1*$^{-/-}$ cells compared to *Dicer1*$^{-/-}$ cells 5 days after *Dicer1* deletion (d5). The results are plotted as percent input (error bars are standard error).

(D) Chromosomal defects are reduced in *Dicer1*$^{-/-}$ day 5 cells treated with siRNAs (n > 100, * - p-value between 0.01 and 0.05, Fisher's exact test).

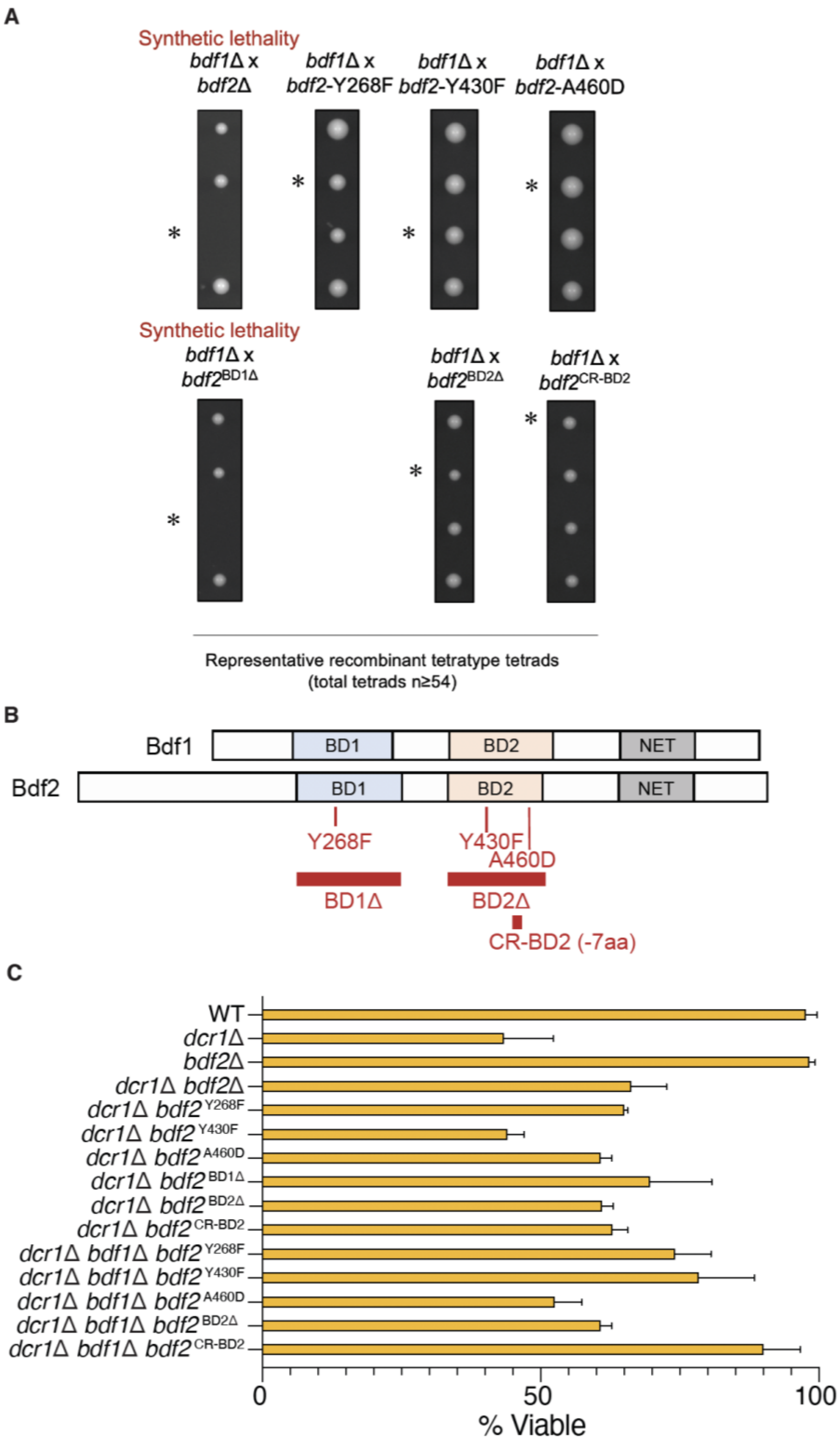


Figure S10. *bdf1Δ* and *bdf2Δ* in *S. pombe* are synthetic lethal via bromodomain 1 (BD1), but suppress *dcr1Δ* via BD2.

(A) Representative tetrads and quantification of viabilities in a *bdf1*Δ x *bdf2*Δ cross. The *bdf1*Δ*bdf2*Δ double-mutant is inviable, as determined from analysis of 54 informative tetrads. Likewise, the *bdf1*Δ*bdf2*$^{BD1\Delta}$ mutant is inviable, while other combinations of mutants were fully viable and displayed no phenotype. A star (*) represents the double-mutant from representative "non-parental tetratype" tetrads.

(B) Schematic of Bdf2 mutants used in the tetrad analysis experiments.

(C) Viability at $G_0$-entry (24h) in wild-type, *dcr1*Δ, and viable double and triple-mutants combinations of the Bdf1/Bdf2$^{BRD4}$ genes. *dcr1*Δ*bdf2*$^{BD2\Delta}$ and *dcr1*Δ*bdf2*$^{CRBD2}$ displayed suppression of the loss of viability of *dcr1*Δ similarly to that of *dcr1*Δ*bdf2*Δ. The strongest suppressor is the triple-mutant *dcr1*Δ*bdf1*Δ*bdf2*$^{CRB2}$, which displayed wild-type viability.
